# Random mutagenesis of influenza hemagglutinin identifies new sites which modulate its acid-stability and cleavability

**DOI:** 10.1101/2025.07.25.666873

**Authors:** S. Rimaux, M. F. Oliva, C. Mestdagh, R. Van Berwaer, J. Chen, G. Schoofs, B. Vanmechelen, J. Stroobants, M. Jacquemyn, M. Laporte, P. Maes, K. Vermeire, K. Das, A. Stevaert, L. Naesens

## Abstract

The structural instability of influenza hemagglutinin (HA) is related to its function in low pH-mediated membrane fusion, which requires prior cleavage of the premature HA0 by a host protease. The precise determinants underlying the stability and cleavability of HA remain to be fully understood and have implications for risk assessment of zoonotic influenza A viruses (IAV), viral transmissibility and vaccine production. To address this, we conducted random mutagenesis on early 2009 pandemic H1 HA, followed by selection of acid-stable viruses and detailed profiling of the mutant HAs. This resulted in identification of four mutations, which increase the acid-stability and decrease the fusion-promoting activity of H1 HA, without compromising viral entry and replication in cells. The newly recognized mutations are situated in the globular head, vestigial esterase and membrane-proximal part of H1 HA, in regions involved in the refolding of HA at low pH. A fifth mutation, D346N, is located in the cleavage loop and renders H1 HA0 12-fold resistant to trypsin activation, whereas its cleavage by transmembrane serine protease 2 (TMPRSS2) is not affected. Along this line, we found that the poor cleavage of H16 HA0, which is unusual in carrying an N346 residue, only applies when it is performed by extracellular proteases. Since H16 HA also exhibits a very low fusion pH, we propose that gull H16N3 virus may carry a much more stable HA than other avian IAVs. Collectively, our mutagenesis approach revealed new determinants of HA stability and cleavability, with relevance for viral surveillance and vaccine production.

**IMPORTANCE:** The presence of influenza A viruses (IAV) throughout the animal world, particularly avian species, represents a constant threat for zoonotic infections or a new influenza pandemic. To be transmissible among humans, a zoonotic IAV requires mutations in the viral hemagglutinin (HA) that increase the acid-stability of this mediator of viral entry. Understanding the determinants of HA stability is also important to produce vaccines with high shelf-stability. By combining random mutagenesis with selection of acid-stable viruses, we identified new stabilizing mutations located in different parts of HA. Besides, we discovered a mutation that renders HA resistant to cleavage by extracellular proteases. Since this residue is naturally occurring in H16 HA, we propose that the gull H16N3 virus may differ from other avian IAVs in carrying an environmentally stable HA. Hence, our study delivers new insight in factors that modulate HA acid-stability and cleavability, with relevance for viral surveillance and vaccine production.

## INTRODUCTION

The propensity of influenza A virus (IAV) to cause zoonotic infections and sporadic pandemics in humans is related to the wide diversity of IAV subtypes and clades that circulate in the animal world, particularly among avian species (1). The last influenza pandemic occurred in 2009 and resulted from a swine-origin reassortant virus [A(H1N1)pdm09] that evolved into an endemic and annually recurring seasonal IAV (2). This pandemic spurred significant interest in understanding the molecular determinants of zoonotic IAV adaptation to humans. Among the different viral factors involved (3), the hemagglutinin (HA) is a key player since it controls viral entry into the host cell. This process starts with viral binding to sialylated glycan receptors and uptake by endocytosis. The endosomal pH (∼5) triggers irreversible refolding of HA, resulting in the extrusion of its fusion peptide and fusion of the endosomal and viral membranes, enabling transport of the viral genome segments into the cytoplasm. The protein structures of pre- and post-fusion HA are dramatically different and have been characterized in much detail (4, 5).

To acquire affinity for the upper part of the human respiratory tract, an avian IAV requires HA mutations that switch its tropism from α2,3-to α2,6-linked sialylated glycans (6). Other adaptation mechanisms are less defined and relate to the ability of HA to undergo cleavage activation by human airway proteases (7–10) and, once cleaved, remain stable upon virus shedding in the environment (11). In virus-infected cells, the HA protein is initially synthesized as its relatively stable HA0 precursor, which requires cleavage into the HA1/HA2 pre-fusion form that is fusion-competent yet metastable. A zoonotic HA may plausibly require certain mutations to improve its cleavage efficiency by a suitable protease of the human airways, such as transmembrane serine protease 2 (TMPRSS2) (12). So far, we lack any knowledge about this kind of adaptation process. There is also no understanding of what determines the cleavability of different HA0 subtypes by trypsin, TMPRSS2 or other proteases (8), or how the location (i.e. intra- or extracellular) of HA0 cleavage during viral replication may help to preserve HA in the acid-labile pre-fusion state.

The subtle link between HA stability and viral transmissibility was first observed in ferret studies with an avian H5N1 virus (13) or an engineered A(H1N1)pdm09 virus bearing avian H5 HA (14). In both studies, the airborne-transmitted virus contained an HA mutation that was shown to increase the acid- and thermostability of HA (14, 15). Conversely, reduced transmission in ferrets was noted with a mutant A(H1N1)pdm09 virus that carried a less acid-stable HA than the wild type (WT) virus (16). The association between HA stability and virus transmission is supported by surveillance data in humans since, within three years after its emergence, the A(H1N1)pdm09 virus acquired HA mutations that increased the protein’s stability (16–18). Several of these mutations also proved important for vaccine production, as one mutation conferred superior viral growth characteristics (19), while other stabilizing mutations improved the thermal stability and shelf-life of the vaccines (17, 19, 20).

In the present study, we applied a broad and agnostic random mutagenesis approach to identify novel sites in HA that modulate its acid-stability and cleavability. We chose the HA of an early A(H1N1)pdm09 isolate as our model protein, since its relatively high fusion pH (i.e. low acid-stability) makes it amenable to acquisition of stabilizing mutations. After selecting acid-stable viruses and identifying four mutations associated with a lower fusion pH, we thoroughly investigated their impact on the efficiency of HA-mediated membrane fusion and on the processes of viral entry and replication. A fifth mutation that we identified, D346N, was observed to significantly impair the trypsin-cleavability of H1 HA0. We presumed an analogy with the trypsin-resistant H16 HA0 (8, 21), which differs from most other subtypes in carrying an asparagine (N) at residue 346, and we therefore included H16 HA for this aspect of the study. Finally, we interpreted our biological findings with respect to the protein structure of uncleaved or cleaved HA. By offering new information on the determinants of HA stability and cleavability, our findings contribute to a better understanding of this protein’s biological properties and its role in IAV species adaptation.

## MATERIALS AND METHODS

### Cells and media

Madin-Darby canine kidney (MDCK) cells were kindly provided by M. Matrosovich (Marburg, Germany). MDCK^TMPRSS2^ cells, a generous gift from J. Bloom (Seattle, USA), are a stable TMPRSS2-tranfectant line that was generated (22) from MDCK-SIAT1 cells, which overexpress 2,6-sialyltransferase (23). Both cell lines were maintained in Dulbecco’s modified Eagle’s medium (DMEM) supplemented with 10% fetal calf serum (FCS), 1 mM sodium pyruvate and 750 mg/L sodium bicarbonate. A similar medium with 10% FCS was used to grow HEK293T (Thermo Fisher Scientific #HCL4517) and HeLa cells (ATCC #CCL-2). The two HeLa-based split-GFP cell lines were prepared as described (24, 25). Briefly, lentiviruses were produced by transfecting HEK293T cells with 10 µg of the pCL-10A1 retrovirus packaging vector (Novus Biologicals), together with 5 µg of either pQCXIP-GFP1-10 or pQCXIP-BSR-GFP11 plasmid (received from Y. Hata through Addgene plasmids # 68715 and #68716), using TransIT-LT1 Transfection Reagent (Mirus Bio). Lentiviruses were harvested at 48 and 72 h post-transfection. Next, HeLa cells were seeded at 500,000 cells per well in 6-well plates and transduced with 600 µL lentivirus together with 10 µg of polybrene, followed by puromycin selection one day later. The resulting polyclonal HeLa-GFP1-10 and HeLa-GFP11 cells were cultured in medium supplemented with 0.6 µg/mL puromycin. All cell lines were kept in a humidified 5% CO_2_ incubator at 37 °C.

### Plasmids

For reverse genetics of A/Puerto Rico/8/34 (PR8)-based chimeric IAVs, we used the pVP-PB2, -PB1, -PA, -NP, -M and -NS plasmids encoding the internal genes of PR8, which were kindly provided by M. Kim (Korea Research Institute of Chemical Technology). In the pVP-HA plasmid, the PR8 sequence was replaced by that of A/Virginia/ATCC3/2009 (A(H1N1)pdm09) [ATCC VR-1738; abbreviated Virg09], by swapping either the entire coding sequence (= Virg09_full_-HA) or only the ectodomain (= Virg09_ecto_-HA). In the pVP-NA plasmid, we replaced the entire NA-coding sequence of PR8 with that of Virg09. The H16 HA and N3 NA sequences of A/black-headed gull/Sweden/5/1999 (abbreviated Gull99) were ordered as GeneArt DNA strings from Life Technologies. To replace the sequences, high-fidelity PCR (Platinum SuperFi PCR, Invitrogen) was performed on the PR8 pVP-plasmids, and Virg09-HA and -NA plasmids prepared earlier (26), using overlapping primers. The amplified Virg09 and ordered Gull99 DNA fragments were inserted into the linearized pVP-plasmid (NEBuilder HiFi DNA Assembly Cloning Kit, New England Biolabs). To create pCAGEN expression plasmids encoding these Virg09 and Gull99 HA and NA sequences, we used high-fidelity PCR and primers extended with EcoRV and NotI sites, to allow subcloning into the pCAGEN vector [provided by C. Cepko (Boston, MA) via Addgene (plasmid 11160)] (27). Specific mutations were introduced into the pVP or pCAGEN plasmids, via site-directed mutagenesis with mutagenic primers and Platinum SuperFi II DNA Polymerase (Invitrogen).

### Reverse genetics

The eight-plasmid reverse genetics procedure was adapted from Martínez-Sobrido and García-Sastre (28) and has been described in detail before (29). To engineer the chimeric Virg09_PR8_ virus, a co-suspension of HEK293T and MDCK cells was prepared in medium with 10% FCS, transferred to a 12-well plate at 1 million cells per well of each cell line, and transfected with 4 µL Lipofectamine 2000 (Thermo Fisher Scientific) and 0.5 µg of each of the eight pVP-plasmids per well. On the next day, the medium was replaced by medium with 0.3% bovine serum albumin (BSA) and 5 µg/mL tosylphenylalanylchloromethylketon (TPCK)-treated trypsin (Sigma-Aldrich) and the plates were then incubated at 35 °C. Two days later, the supernatant was collected to expand the virus in MDCK cells, using infection medium [i.e. UltraMDCK (Lonza) supplemented with 225 mg/L sodium bicarbonate, 2 mM L-glutamine, 100 U/mL penicillin-streptomycin and 5 µg/mL TPCK-treated trypsin]. The virus was collected at day 3 post-infection (p.i.) and stored in aliquots at −80 °C. Titration was done by end-point dilution in 96-well plates containing 7,500 MDCK cells per well. After scoring the cytopathic effect (CPE) at day 3 p.i., the 50% cell culture infective dose (CCID_50_) was calculated using the method of Reed and Muench (30).

To successfully rescue the chimeric Gull99_PR8_ virus, the above method required some modifications, i.e. transfection of a co-suspension of HEK293T and MDCK^TMPRSS2^ cells, and addition of a TMPRSS2 expression plasmid (26) to the eight pVP plasmids in the transfection mix.

To determine viral growth kinetics, the virus was inoculated on MDCK or MDCK^TMPRSS2^ cells at a multiplicity of infection (MOI) of 200 CCID_50_ per well. The infection medium (see above) contained 5 µg/mL trypsin (for MDCK) or no trypsin (for MDCK^TMPRSS2^). The supernatants were harvested at 24, 36, 48 and 72 h p.i., then titrated in MDCK cells.

### Generation of randomly mutated virus libraries and selection of acid-stable HA-mutants

To create virus libraries with random mutations in HA, we performed error-prone PCR (GeneMorph II EZClone Domain Mutagenesis Kit, Agilent) on the Virg09_ecto_-HA sequence in the pVP-plasmid. Two primer sets were used (sequences in Table S1) to separately amplify two parts of the HA sequence. The PCR products served as megaprimers for the EZClone reaction. Based on a concise optimization experiment, we selected the condition of 500 ng input DNA and 30 PCR cycles, to obtain a mutation frequency of approximately one mutation per sequence. In this way, six randomly mutated pVP-Virg09_ecto_-HA libraries were obtained (= two fragments submitted to three separate PCR reactions). These were used to conduct reverse genetics as outlined above, and produce six libraries of HA-mutant viruses.

To select acid-stable mutants, the undiluted virus libraries were adjusted to pH 5.0 by adding a predetermined amount of citric acid, then incubated for 1 h at 37 °C. After a 1:4 dilution in neutral MDCK infection medium (to reduce the acidity), the viruses were added to MDCK cells seeded at 43,000 cells per well in 24-well plates, followed by medium replacement at 1 h p.i. After three days incubation, the wells showing CPE were selected and these supernatants were harvested. Following RNA extraction (QIAamp Viral RNA Mini Kit, Qiagen), the SuperScript One-Step RT-PCR System from Invitrogen was used to amplify the full HA sequence with two primer pairs: the H1R1264-H1F848 pair recommended by the WHO (https://www.who.int/teams/global-influenza-programme/laboratory-network/quality-assurance/eqa-project/information-for-molecular-diagnosis-of-influenza-virus), and another pair targeting the PR8 5′ and 3′ ends of the chimeric HA sequence. Finally, the HA fragments were submitted for Sanger sequencing by Macrogen Europe.

### Cell-cell fusion assays in HA-expressing cells, based on luminescence, split-GFP or impedance

#### Luminescence

To obtain a quantitative readout for HA-mediated cell-cell fusion, we used the pGal5-luciferase and pGal4-VP16 plasmids (kind gift from S. Pöhlmann, Göttingen, Germany) and transactivation setup reported by his team (31), with several modifications. To minimize luminescence cross-over, the assay was conducted in white 96-well plates. On day 0, the HeLa target cells were reverse-transfected at a ratio (expressed per well) of 15,000 cells, 50 ng HA-pCAGEN plasmid, 13 ng pGal4-VP16 plasmid and 0.26 µL Fugene 6 (Promega). In parallel, a 6-well plate was prepared with, in each well, 2 million HeLa effector cells, 500 ng pGal5-luciferase plasmid and 2 µL Fugene 6. On the next day, the effector cells were detached using non-enzymatic cell dissociation solution (Sigma-Aldrich), resuspended and overlaid on the target cells at 10,000 cells per well. After another 24 h, the medium was replaced with blank medium, and HA was activated by 15 min incubation with 5 µg/mL TPCK-treated trypsin. Next, cell-cell fusion was induced by incubation in acidic buffer (= PBS with Ca^2+^ and Mg^2+^, adjusted to pH with acetic acid) for exactly 5 min. To apply different pH conditions across wells, a series of buffers ranging from pH 5.0-6.0 in 0.1 increments was used. After replacing the acidic buffer with growth medium, the plates were incubated for five hours at 37 °C. Finally, the cells were lysed and luminescence was measured using the Luciferase Assay System and Glomax Navigator (both from Promega), using injector mode and 100 µL buffer per well. The percentage fusion was calculated by subtracting the background signal at pH 7 from the signal at each pH tested, then dividing this value by the one obtained at pH 5.0.

#### Split-GFP

The fluorescent assay was conducted in black-wall 96-well plates. Reverse transfection was performed at a ratio (expressed per well) of 10,000 HeLa-GFP1-10 cells, 10,000 HeLa-GFP11 cells, 50 ng HA-pCAGEN plasmid and 0.26 µL Fugene 6. Two days later, HA activation and induction of cell-cell fusion were done as described above for the luminescence assay. After another 24 h, fluorescent focusing beads (Bangs Laboratories) were added at 1,250 beads per well, and the plates were placed in a CellInsight CX5 high-content imaging instrument to quantify the total GFP area via the SpotDetector protocol of the imaging software. The percentage fusion was calculated by subtracting the background value at pH 7 from the GFP area at each pH tested (4.5-5.5 for H16 HA and 5.0-6.0 for H1 HA), and dividing this value by the GFP area at pH 5.0 (H1 HA) or 4.5 (H16 HA).

#### Impedance

The impedance protocol was adapted from (32) and conducted in 16-well plates with gold microelectrodes (Agilent). On day 0, HeLa cells were reverse-transfected at a ratio of 20,000 cells, 50 ng HA-pCAGEN plasmid and 0.26 µL Fugene 6 per well, after which the plate was placed in an xCELLigence instrument (Thermo Scientific). At 48 h post-transfection, a short baseline measurement of the cell index (CI) was performed (5 consecutive measurements, every 5 s). Next, the HA was activated with 5 µg/mL trypsin, after which cell-cell fusion was induced by 10 min incubation in pH 5.0. The formation of syncytia was followed in real-time, by recording the CI value for 20 h at 2-min intervals. Data were analyzed as in (32), using the Matlab script (version R2016b, Mathworks) to normalize the raw CI values to the baseline value, followed by Graphpad Prism 10.3.1 to calculate the area under the curve (AUC).

### Pseudovirus entry assay

To produce murine leukemia virus (MLV)-based pseudoviruses bearing the HA and NA of interest (26), HEK293T cells were seeded in a 6-well plate at 700,000 cells per well, in medium containing 0.2% FCS and 0.3% BSA. On the next day, they were transfected at a ratio (expressed per well) of 6 µL Lipofectamine 2000, 750 ng MLV-packaging vector and 1500 ng luciferase plasmid (both kindly donated by S. Pöhlmann), combined with 1000 ng pCAGEN-HA plasmid and 250 ng pCAGEN-NA plasmid. At day 3 post-transfection, the HA-bearing pseudovirus was activated by adding TPCK-treated trypsin at a final concentration of 80 µg/mL (unless specified otherwise). After 15 min incubation at 37 °C, the same concentration of soybean trypsin inhibitor (Sigma-Aldrich) was added. In experiments assessing HA0 cleavability by TMPRSS2, −4 or −13, the pseudovirus was not activated by trypsin but, instead, 270 ng (unless specified otherwise) of a plasmid encoding TMPRSS2, −4 or −13 (26) was combined with the other four plasmids on day 0 of the production scheme. All pseudovirus stocks were stored in aliquots at −80 °C.

To determine the pseudovirus entry capacity, 3,750 MDCK or MDCK^TMPRSS2^ cells were seeded in white half-area 96-well plates, in medium containing 0.2% FCS and 0.3% BSA. On the next day, the medium was replaced by the same medium supplemented with 10 µM zanamivir (Sigma-Aldrich). Subsequently, the pseudovirus was added and entry was promoted by spinoculation for 45 min at 450 x g and 37 °C, followed by 60 min in the incubator at 37 °C. Next, the supernatant was replaced by fresh medium, and the plates were incubated for three days at 37 °C. Expression of the firefly luciferase reporter was quantified using a Glomax Navigator and luciferase assay system, as described above.

### Assessment of HA0 cleavage

To assess the extent of HA0 cleavage by trypsin or TMPRSS2, a 600-µL aliquot of the pseudovirus stock was loaded on 50 µL of 20% sucrose in PBS, and centrifuged for 2 h at 21,000 x g and 4 °C. The bottom fraction was collected and lysed in RIPA buffer supplemented with protease inhibitor cocktail and EDTA (both from Thermo Fisher Scientific). The lysates were treated with PNGase F, as instructed by the manufacturer (New England Biolabs), to deglycosylate the HA0, HA1 and HA2 proteins and optimize their separation in the Simple Western analysis. Capillary electrophoresis, antibody binding and detection were performed according to the manufacturer’s instructions. All steps were done using default settings, except for the separation step which was set to 35 min. Specifically, proteins were size-separated on the 12-230 kDa Jess Separation Module (SMW004) and bound with primary anti-HA antibody [i.e. rabbit anti-H1 HA (SinoBiological 11055-RM05; 1/25 diluted) or goat anti-H16 HA (BEI resources NR-34798; 1/40 diluted)], followed by secondary antibodies of detection modules DM-001 and DM-006. Next, the primary and secondary antibodies were removed using the Replex Module (RP001), to allow subsequent total protein quantification. Finally, the protein signals were visualized via Compass for Simple Western software, v.6.1.0 (ProteinSimple). To quantify the percentage cleavage, the corrected peak areas of HA1 or HA2 were divided by the sum of the corrected peak areas of HA0 plus HA1 or HA2.

### Flow cytometric analysis of HA and TMPRSS2 expression

HeLa cells were reverse-transfected with HA in the same way as in the cell-cell fusion assays, and seeded in a 6-well plate. Two days later, the cells were washed with blank medium, then incubated with 5 µg/mL TPCK-treated trypsin to activate HA. Next, they were detached with non-enzymatic cell dissociation solution and resuspended in medium with 10% FCS, after which they were allowed to regenerate for 1 h. Then, the cells were pelleted, resuspended in PBS and stained with BD Horizon™ Fixable Viability Stain 780. A cell death control was prepared by heating at 65 °C for 5 min. Subsequently, the cells were placed on ice and stained for 30 min with primary antibody [= mouse anti-HA antibody C179 (Takara M145; 10 µg/mL)], followed by 30 min incubation with secondary antibody [anti-mouse IgG AF488 (Invitrogen A21131; 5 µg/mL)], with washing steps in between. All stainings were done in FACS buffer (= PBS with 2% FCS). In the last step, the cells were fixed in 2% paraformaldehyde for 5 min. Data were acquired on a FACS Celesta flow cytometer (Becton Dickinson) and analyzed with FlowJo software.

To measure expression of TMPRSS2, HEK293T cells were transfected with the TMPRSS2 plasmid, in the same way as done for the pseudovirus production. At 72 h post-transfection, the cells were detached with non-enzymatic cell dissociation solution and resuspended in medium with 10% FCS, in which they were allowed to regenerate for 1 h. Afterwards, the staining procedure and flow cytometric analysis were conducted as described above. The primary antibody was rabbit anti-TMPRSS2 antibody from Abcam (ab280567; 0.5 µg/mL) and the secondary antibody was anti-rabbit IgG from Cell Signaling Technology (4414S; 1 µg/mL).

### HA models and graphics

To study the stabilizing mutations and cleavage loop structures on a molecular level, SWISS-MODEL (33) was used to model our strain sequence on several cryo-EM and crystal structures. AlphaFold was used for modeling of the cleavage loops (34), and ChimeraX was used to make graphics (35).

### Schematics, data analysis and statistics

Graphical schemes were created using BioRender (Naesens, L. (2025) https://BioRender.com/z9ws7tt) and edited in PowerPoint and Photoshop. All graphs were created with GraphPad Prism 10.3.1 software, which was also used for statistical analysis. Statistical significance is shown as: \*\*\*\**P* ≤ 0.0001, \*\*\**P* ≤ 0.001, \*\**P* ≤ 0.01, \**P* ≤ 0.05, ns: not significant.

## RESULTS

### Engineering of chimeric H1N1 virus with early 2009 pandemic HA sequence

To identify mutations that render the HA protein more acid-stable, we required an HA that was relatively unstable. We therefore selected the H1 HA from strain A/Virginia/ATCC3/2009 (abbreviated Virg09), which was isolated early after the start of the H1N1 pandemic in 2009. The swine-origin A(H1N1)pdm09 virus initially carried an HA with intermediate stability (fusion pH ∼5.5). Upon sustained circulation in the human population, this virus acquired HA mutations associated with increased acid-stability and transmissibility (16–18).

The HA and NA sequences of Virg09 were used to reverse-engineer a chimeric virus that carried the six internal genes of A/Puerto Rico/8/1934 (PR8) and was therefore referred to as Virg09_PR8_. To optimize replication fitness, we initially compared two approaches (36): (i) replacing the full-length PR8 HA with that of Virg09 (Virg09_full_-HA) or (ii) retaining the signal peptide, transmembrane and cytosolic tail domains of PR8 HA while introducing only the ectodomain of Virg09 HA (Virg09_ecto_-HA). In both cases, the entire PR8 NA coding sequence was replaced with that of Virg09, while preserving the PR8 non-coding regions. Consistent with a previous report (36), the Virg09_ecto_-HA approach yielded slightly higher virus titers (data not shown), although the difference was not significant.

### Random mutagenesis and selection of HA mutations conferring higher acid-stability

To perform random mutagenesis on the Virg09_ecto_-HA plasmid, we conducted error-prone PCR using Mutazyme II polymerase (Fig. 1). Since the mutation rate is easier to control on smaller DNA fragments, we split the HA sequence into two parts and used two primer sets for amplification. A low mutation frequency was ensured by following the manufacturer’s instructions and conducting a pilot experiment under varying conditions (i.e. 100 or 500 ng target DNA and 20 or 30 PCR cycles). Sequence analysis of plasmids from twenty transformed *E. coli* colonies showed that most contained a single mutation in the HA gene when using 500 ng DNA and 30 cycles. In this way, we generated six mutagenized HA plasmid libraries (i.e., two HA parts and three separate PCR reactions), which served to reverse-engineer six libraries of Virg09_PR8_ virus carrying random HA mutations.

**FIGURE 1.**
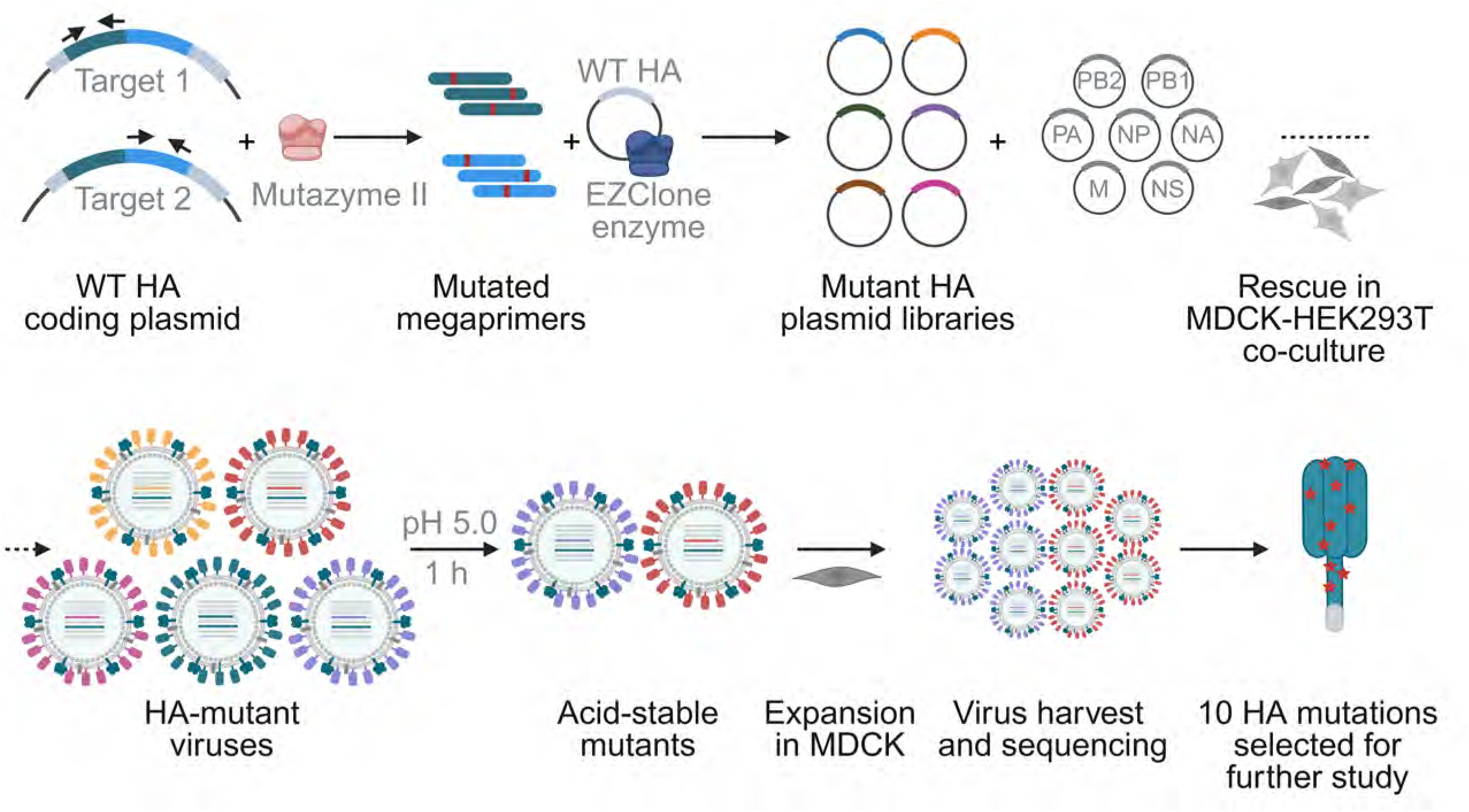
Procedure for random mutagenesis and selection of HA-mutant viruses with higher acid-stability. The reverse genetics plasmid encoding Virg09_ecto_-HA was submitted to error-prone PCR and combined with the other seven plasmids to generate HA-mutant libraries of Virg09_PR8_ virus. After 1 h incubation at pH 5.0, the surviving mutants were expanded and sequenced. This yielded ten HA mutations for follow-up.

To select mutant viruses with a more acid-stable form of Virg09 HA, the Virg09_PR8_ libraries were incubated for 1 h at pH 5.0 (37 °C), then applied to MDCK cells to expand viruses that survived the low pH treatment (Fig. 1). Supernatants from wells showing viral cytopathic effect (CPE) were harvested for HA sequencing. [Note: throughout this report, we use the same H1 HA0 numbering as in another study on HA-stabilizing mutations (17); see Table S2 for the corresponding H3-numbering]. Only one of the selected viruses carried more than one HA mutation, confirming the low mutation frequency of the PCR. All ten mutations were individually introduced in an HA expression plasmid to assess their impact on HA fusion pH using a luciferase-based cell-cell fusion assay (Fig. 2A). For wild-type (WT) Virg09_ecto_-HA, we measured a fusion pH of 5.43 (Fig. 2B). This value aligns with our previous measurement for full-length Virg09 HA (26), and with the fusion pH of 5.5 reported for an early A(H1N1)pdm09 isolate (16). Within three years after the 2009 pandemic, 100% of circulating isolates carried HA mutation E374K (= E47_2_K in H3-numbering; see Table S2) a key marker for the virus’s rapid evolution toward higher HA stability (17, 19). We therefore included mutation E374K as reference and found its fusion pH to be 5.17 (Fig. 2B), which is 0.26 pH units lower than our WT (*P* < 0.0001). After analyzing the pH profiles and calculating the corresponding fusion pH values, we identified three mutations (i.e. I79V, T241S and K454N) that significantly increased acid-stability (*P* < 0.001 or *P* < 0.0001 *versus* WT). A fourth mutation, K211M, caused a small, non-significant decrease of 0.05 pH units but was still retained for further study to explore whether alternative substitutions at this position could lead to a more pronounced reduction in fusion pH. When we engineered the K211T mutant instead, a significant reduction in fusion pH by 0.11 units was observed (*P* < 0.0001 *versus* WT). A few mutations identified after the pH 5.0 selection step did not reduce the fusion pH (Fig. S2) and were therefore omitted from further study.

**FIGURE 2.**
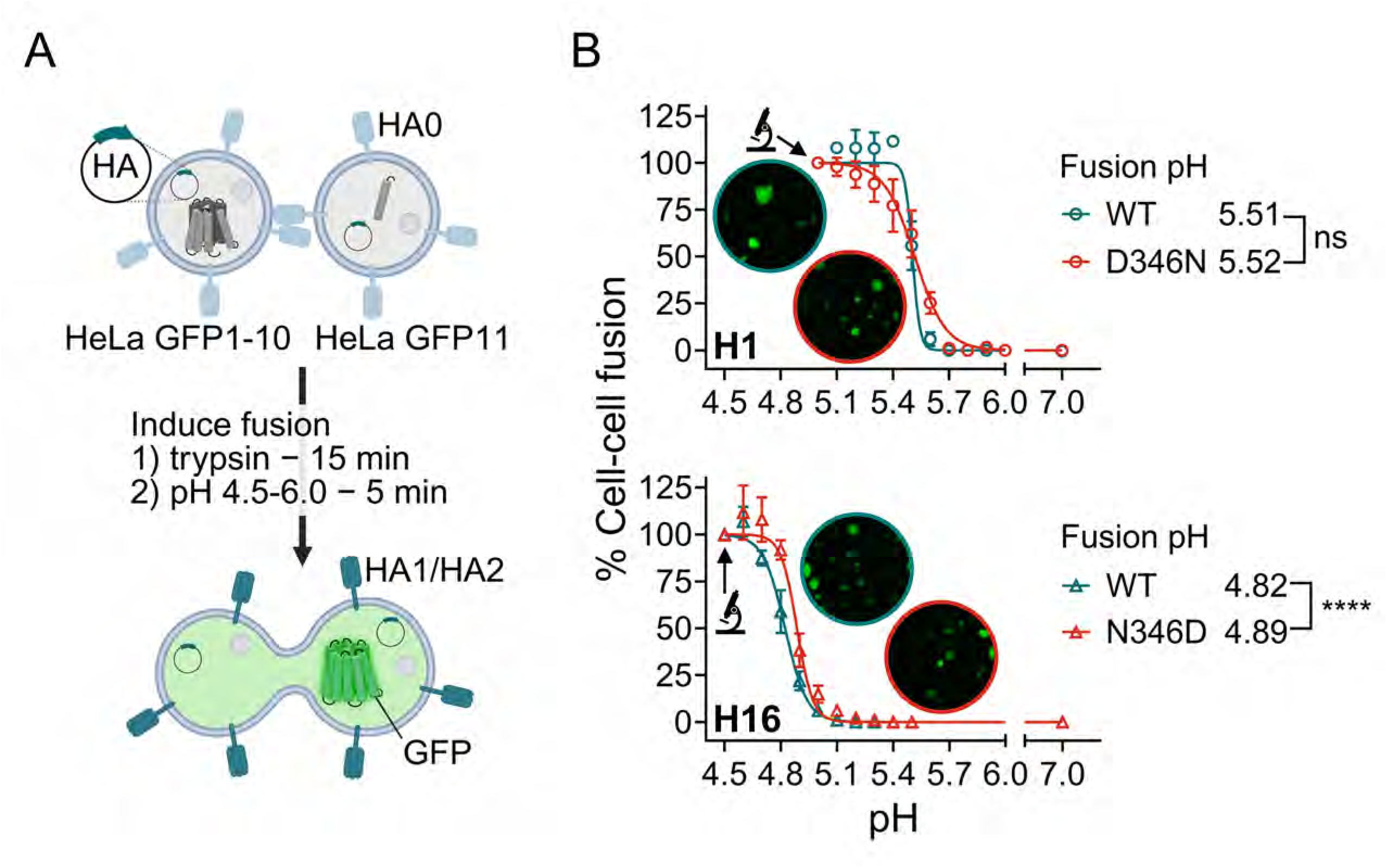
pH profile of the mutations which render H1 HA more acid-stable. (A) Scheme of the luciferase reporter-based cell-cell fusion assay in HA-expressing cells exposed to low pH. (B, C) pH profiles for WT HA (black curve that is shown in each graph) and mutant forms of HA (curves in red). Panel B: reference mutation E374K, four mutants that exhibited a shift in fusion pH and two variations. Panel C: triple (I79V + K211T + K454N), quadruple (= triple + E374K) and quintuple mutant (= quadruple + T241S). In each graph, the Y-axis shows the % cell-cell fusion, calculated by dividing the luminescence signal (after background subtraction) at each pH by the one at pH 5.0. The legend shows the fusion pH, defined as the pH where fusion was 50% relative to pH 5.0. Statistical significance is shown for the difference between mutant and WT (Extra sum of squares F test of best-fit value). Data points are the mean ± SEM (N=3-4).

Given that the five stabilizing mutations are distributed across distinct regions of the HA protein (see below; last section of results), we hypothesized that combining them might yield a cumulative stabilizing effect. Therefore, we evaluated mutant HAs bearing I79V + K211T + K454N (= triple mutant; fusion pH 5.21), combined with E374K (= quadruple mutant; fusion pH 5.12) and T241S (= quintuple mutant; fusion pH 5.16) (Fig. 2B). Hence, the triple mutant proved only slightly more stable than the individual I79V, K211T and K454N single mutants (fusion pH: 5.29, 5.33 and 5.29, respectively; Fig. 2B). Similarly, the quadruple and quintuple mutants barely reached higher stability than the E374K mutation alone (fusion pH: 5.17), suggesting that the combined effect of these mutations is, at best, modestly additive.

Finally, our pH 5.0 selection procedure yielded one HA mutation, D346N, which seemed to abolish fusion capacity under our assay conditions. In both luminescence-based and microscopic cell-cell fusion assays, the D346N-mutant HA failed to induce syncytia. Because of this intriguing finding, we conducted an in-depth investigation of mutation D346N in subsequent experiments.

### Most of the mutated residues show high conservation rate in H1 HA

We next analyzed HA sequences in the Influenza Research Database (38), using the single nucleotide polymorphisms (SNP) tool, to assess whether the selected HA mutations occur naturally in human, swine (pandemic or non-pandemic) or avian H1 sequences. The amino acid variation for the five residues is shown as a heatmap in Fig. 3.

**FIGURE 3.**
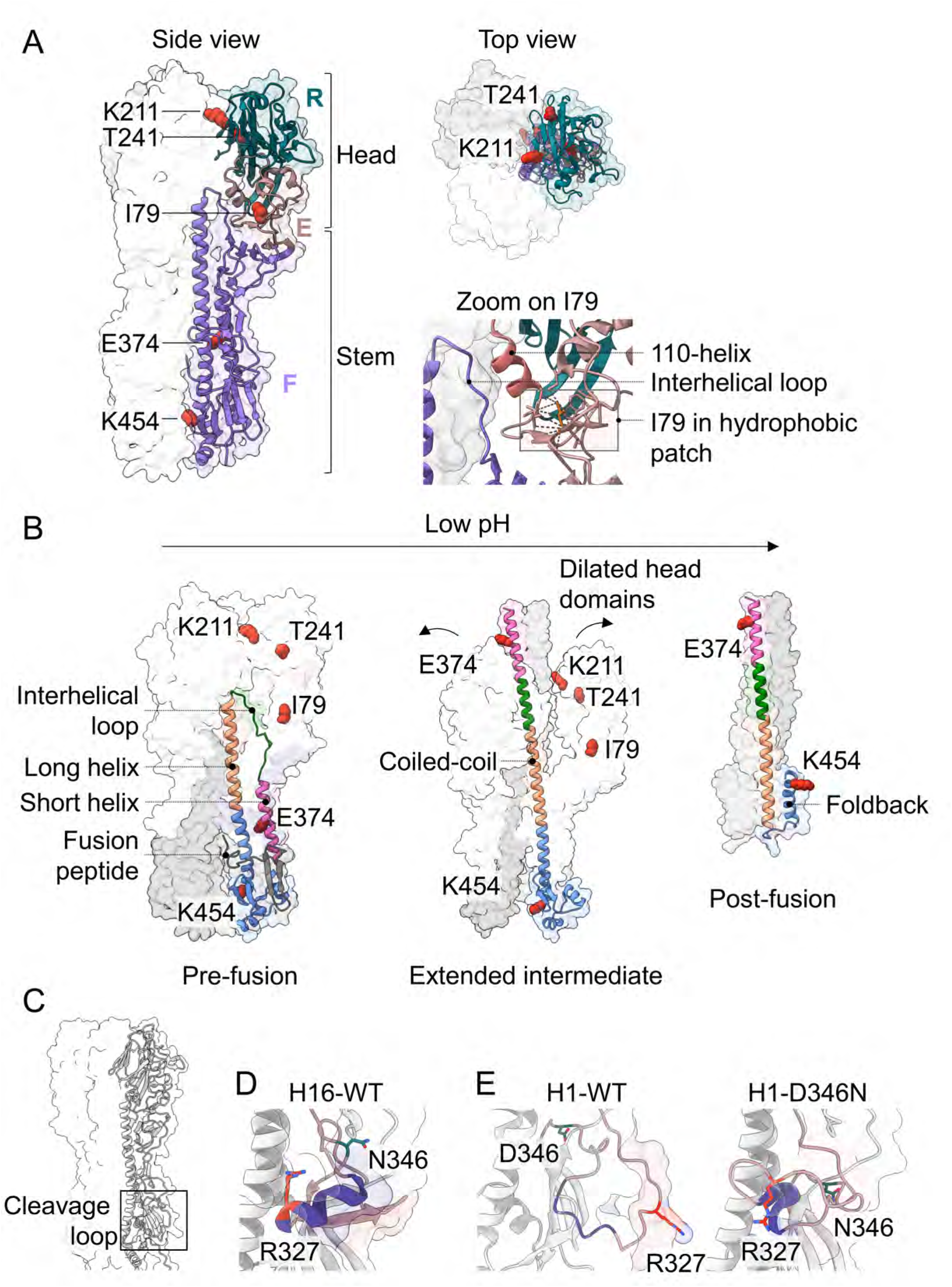
Conservation rate of the acid-stability determining residues in different subgroups of H1 HA. The Influenza Research Database online single nucleotide polymorphisms (SNP) tool was used to determine the % of sequences bearing any of the 20 amino acid residues at the five mutated positions. From left to right and separated by a dotted line: basic, acidic, neutral, hydrophobic and other residues. Note that the number of sequences was not equal in the five subgroups (see graph at the top). The heatmaps visualize the % of the sequences bearing the residues on the X-axis, with the color legend shown on the right.

Overall, four of the five residues showed a fairly high degree of conservation in H1 HA. Residue I79 is present in ∼99% of the ∼40,000 H1 HA sequences in our analysis, and the few variations at this position consistently involve another hydrophobic residue. At site 241, threonine is dominant in H1 HAs, including the avian subgroup. I241 occurs in approximately 2% of the non-pandemic human and swine sequences. Residue D346 was detected in 99.8% of all H1 sequences analyzed, and in 100% of the avian ones. Finally, at site 454, a positively charged residue is present in more than 99% of the sequences, with R454 appearing more abundant in avian H1 HAs and K454 dominating in human and swine sequences. In about 1% of swine H1 HAs, the positive residue is substituted by asparagine; in the other host species, substitutions of R/K454 are extremely rare. Based on this observation, we performed a cell-cell fusion assay with a mutant HA that carries a negatively charged residue at this position, specifically the K454D-mutant of Virg09 HA. Its fusion pH was measured to be 5.32 (Fig. 2B), which was significantly different from WT (*P* < 0.01) and similar to that of mutant K454N (fusion pH 5.29).

The fifth residue, K211, appeared to be the least conserved, since diverse amino acids are tolerated at this site. Variation M211 (picked up from our mutagenesis and pH 5.0 selection approach) proved only rarely present, while T211 is very common and even dominant in the avian reservoir. In non-pandemic swine strains, E211 (which introduces a negatively charged residue) is quite widespread.

Hence, except for K211T, the stabilizing mutations identified in this study involve residues that are highly conserved among circulating H1 HAs, even across human, swine and avian hosts.

### The stabilizing HA mutations reduce the efficiency of cell-cell fusion without affecting the overall process of viral entry

To determine whether the identified mutations affect the fusion-promoting activity of HA, we assessed cell-cell fusion using a real-time, impedance-based assay. This method, previously established in our laboratory for monitoring coronavirus spike-mediated fusion (32), tracks changes in electrical impedance in growing and fusing cells (Fig. 4A). HA-expressing HeLa cells were cultured for 48 hours to form a confluent monolayer, resulting in a gradual increase in cell index (CI). Cells were then treated with trypsin to activate HA, followed by exposure to pH 5.0 to trigger fusion. As cell-cell borders dissolve and syncytia form, electrical resistance increases, resulting in a sharp rise in CI. This is followed by a decline due to syncytial instability and subsequent cell lysis. Fusion efficiency was quantified as the area under the curve (AUC), which correlates with syncytium size (32).

**FIGURE 4.**
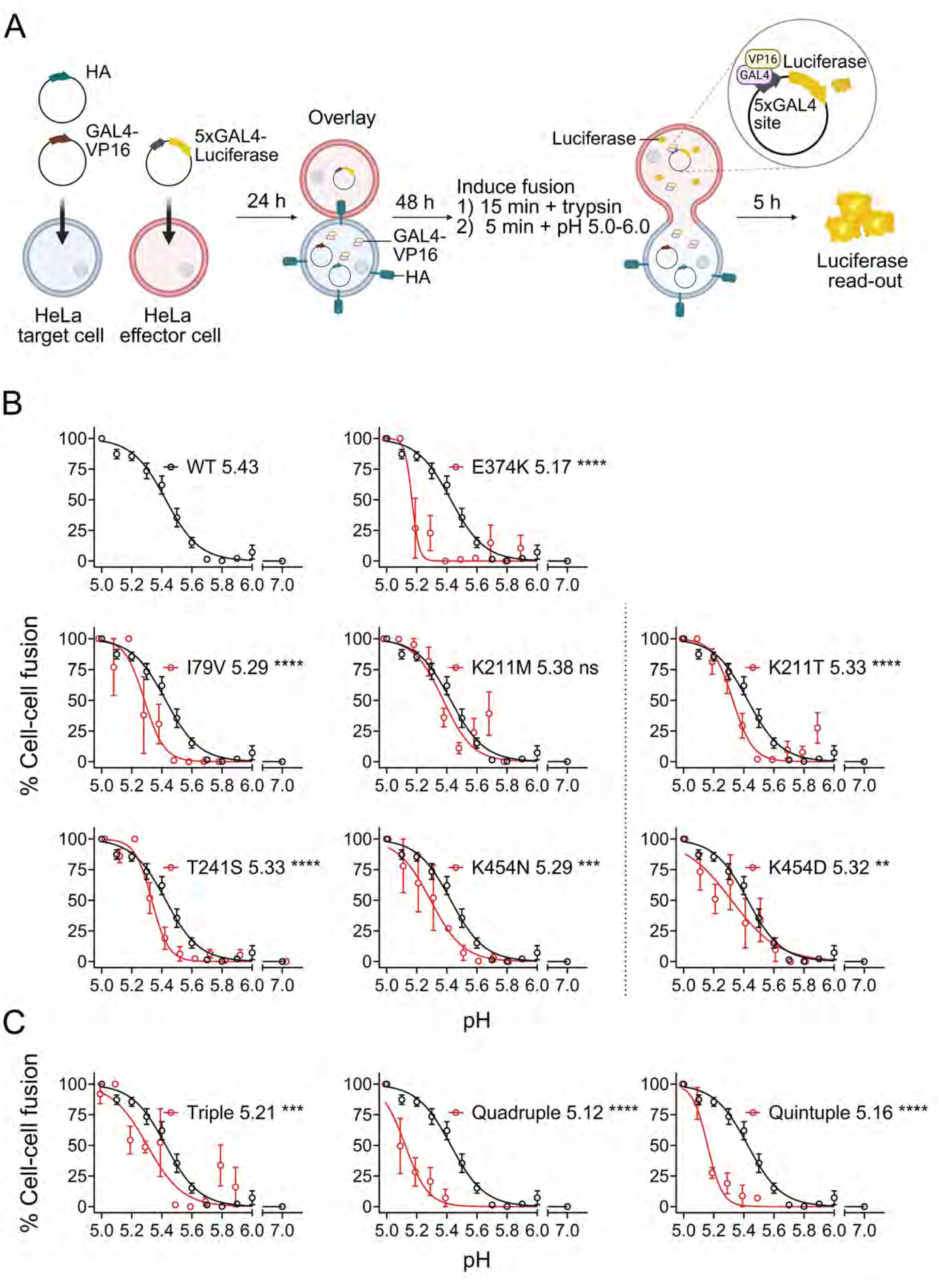
The HA mutations affect the efficiency of cell-cell fusion. (A) Assay scheme. (B) HeLa cells were transfected with WT or mutant HA and, 48 h later, analyzed for HA surface expression using flow cytometry. CC: cell control transfected with empty plasmid. MFI: mean fluorescence intensity; individual and mean values ± SEM (N=3). For neither of the mutants, the MFI was significantly different compared to the WT (not shown on the graph; ordinary one-way ANOVA with Dunnett’s corrections for multiple comparisons). (C, D) In parallel, the cells were sequentially exposed to trypsin and pH 5.0 (= time 0) to induce syncytium formation, which was followed by measuring the impedance during 20 h. Panel C: area under the curve (AUC) for each HA and calculated from the pooled cell index data from 3 (mutants) or 6 (WT, mock and E374K reference) independent experiments. For each mutant except for I79V, the difference versus WT was significant (ordinary one-way ANOVA with Dunnett’s corrections for multiple comparisons). Mock: cells transfected with empty (instead of HA) plasmid. Panel D: graph showing the normalized cell index in function of time. The dotted lines indicate the baseline that was used to normalize the values, which was measured right before cell-cell fusion was induced.

To ensure that differences in fusion were not due to altered HA expression at the cell surface, we next quantified HA surface levels by flow cytometry, conducted in parallel with the impedance assay. As shown in Fig. 4B, all mutants exhibited comparable HA surface expression levels. In sharp contrast, all mutants, including the E374K reference, showed significantly reduced AUC values relative to WT Virg09-HA (*P* < 0.0001 or *P* < 0.001, Fig. 4C), with the exception of I79V. The D346N mutant displayed a nearly flat impedance profile (Fig. 4D), consistent with its complete loss of fusion activity (see above). E374K caused the most pronounced reduction (3.1-fold) in syncytium size, while K211T and T241S led to moderate reductions (1.5-to 2.4-fold). K454N caused a slight (1.1-fold) reduction in AUC compared to WT. Combining I79V with K211T and K454N (= triple mutant) further impaired syncytium formation, and this effect was further exacerbated by the addition of E374K (= quadruple and quintuple mutants) (Fig. 4C and 4D).

To assess whether these mutations affect HA function during viral entry, we generated luciferase-expressing pseudoviruses bearing WT or mutant HA, along with Virg09 NA. Pseudoviruses were treated with trypsin to activate HA, i.e. to cleave the precursor HA0 into HA1 and HA2. These activated pseudoviruses were used to transduce MDCK cells in the presence of 10 µM zanamivir to inhibit NA activity and preserve sialylated receptors. Luciferase activity measured at 72 h post-transduction served as a readout for viral entry (Fig. 5A). Pseudoviruses carrying HA mutation I79V, K211T, T241S, E374K or K454N, showed no significant difference in entry compared to WT (*P* > 0.9). The quadruple and quintuple mutants exhibited slight reductions, with only the quintuple mutant reaching statistical significance (*P* < 0.01, Fig. 5B). In contrast, the D346N pseudovirus showed a >5000-fold reduction in luciferase signal (*P* < 0.0001, Fig. 5C), comparable to the non-activated (no trypsin) WT control, indicating a severe entry defect.

**FIGURE 5.**
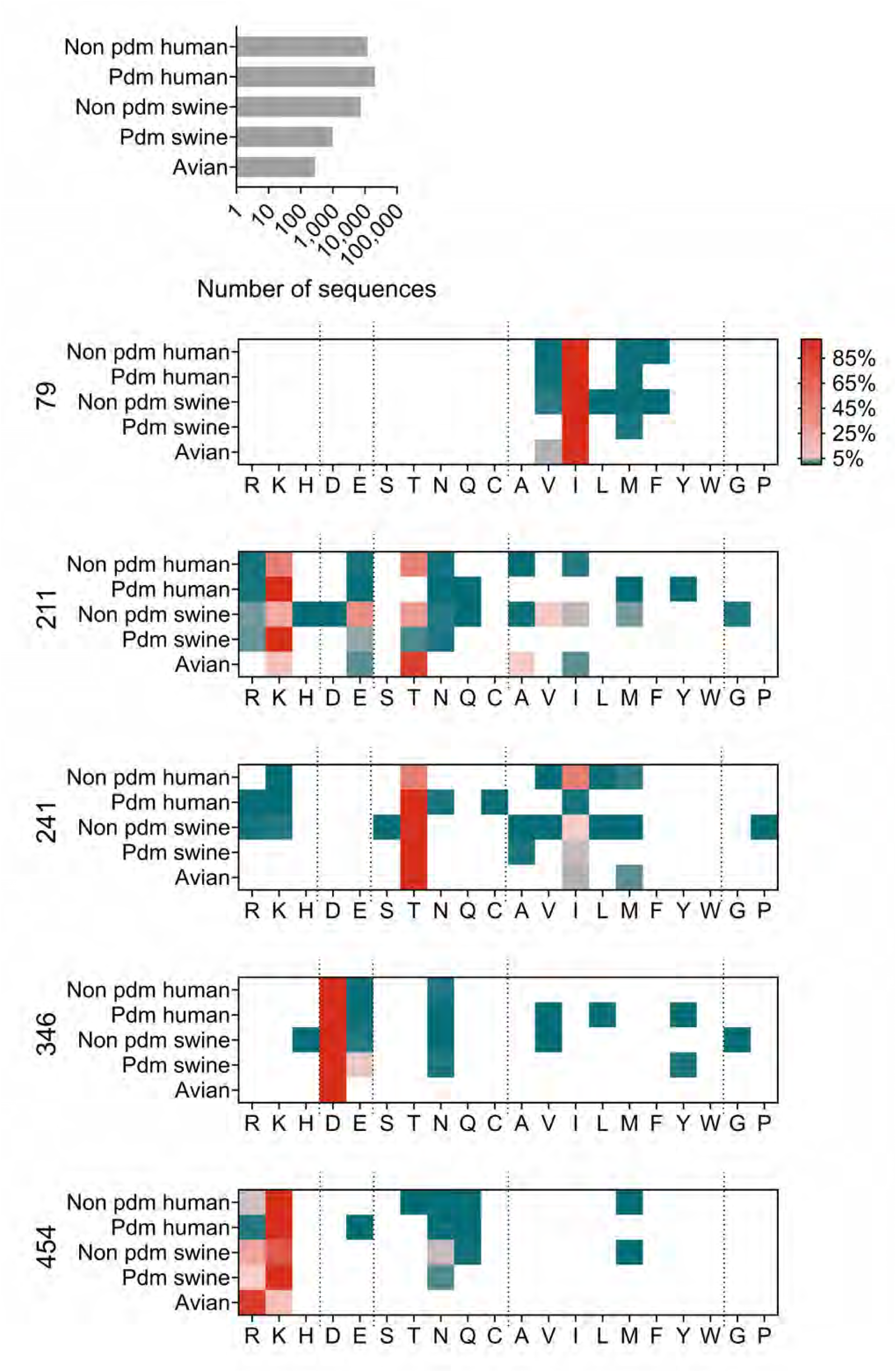
Entry efficiency of HA-mutant pseudoviruses into MDCK cells. (A) After production in HEK293T cells, the HA0-bearing pseudoviruses were activated with 80 µg/mL of trypsin, then incubated with MDCK cells for 72 h, followed by luciferase readout. (B) Relative Luminescence Units (RLU) for mock (= pseudovirus without HA and NA), WT and HA-mutant pseudoviruses. Triple = I79V + K211T + K454N; quadruple = triple + E374K; quintuple = quadruple + T241S. Individual and mean values ± SEM from 3 (mutants) or 6 (WT) independent experiments. Only the quintuple mutant showed significance *versus* WT (ordinary one-way ANOVA with Dunnett’s corrections for multiple comparisons). (C, D) Profile of the D346N mutant. Panel C: RLU values (individual and mean values ± SEM; N=3) in MDCK cells incubated with pseudovirus carrying WT or D346N-mutant HA that was (+) or was not (-) activated with 80 µg/mL of trypsin. Mock: pseudovirus without HA and NA. A two-tailed unpaired t-test was used for statistical analysis. Panel D: HA0 cleavage profile of the four pseudovirus conditions, based on Simple Western analysis.

To evaluate the impact of these mutations on viral replication, we reverse-engineered Virg09_PR8_ viruses carrying the stabilizing HA mutations (single and combined) and passaged them four times in MDCK cells. All viruses replicated to high titers (i.e. 5-6 log_10_ CCID_50_/mL; Table S3) and retained their HA mutation(s) after four passages. This indicates that the stabilizing mutations have little to no negative impact on viral fitness. In some viruses, one to three additional HA mutations emerged, which may have had compensatory effects, as their prevalence declined following low-pH selection (Table S4).

To summarize, our random mutagenesis combined with pH 5.0 selection led to the identification of four residues (I79, K211, T241 and K454) that show moderate to high conservation among H1 HA and were not previously implicated in modulating HA acid-stability. Mutations at these sites significantly lowered the fusion pH and reduced the fusion-promoting activity of HA. However, this does not translate into impairment of viral entry or replication.

In the following part, we focus on a fifth site, D346, where a mutation (D346N) was selected from virus libraries exposed to pH 5.0. Intrigued by the observation that this mutation appeared to abolish the fusion-promoting activity of H1 HA, we sought to investigate the underlying biological mechanism in more detail.

### D346 is conserved in most HA subtypes, but replaced by N346 in avian H11, H13 and H16

Given its location in the HA0 cleavage loop, we hypothesized that the D346N mutation might impair HA activation by interfering with proteolytic cleavage. Supporting this, pseudovirus entry experiments (see Fig. 5C and text above) showed that the D346N mutant yielded a markedly low luciferase signal, comparable to the non-activated (i.e. no trypsin treatment) WT control, suggesting a defect in HA0 cleavage. To directly assess HA activation, we performed Simple Western analysis on pseudovirus particles (Fig. 5D). Trypsin-treated WT pseudovirus showed complete cleavage of HA0, with a prominent HA1 band and no detectable HA0. In contrast, the D346N mutant pseudovirus contained only uncleaved HA0, confirming a cleavage defect.

HA0 cleavage is essential to liberate the fusion peptide, which comprises the N-terminal region of HA2. In precursor HA0 protein, the cleavage loop contains the scissile arginine (= R327 based on the numbering used here), which serves as the recognition site for trypsin-or furin-like proteases in mono- and multibasic HAs, respectively (39). The position of D346 within this loop suggests that substitutions at this site may be poorly tolerated by influenza virus. To explore this further, we aligned the cleavage loop sequences across all 19 IAV-HA subtypes. For each subtype, we aligned 50-1000 sequences per clade to generate clade consensus sequences, which were then used to derive subtype consensus sequences. For H5 clade 2.3.4.4.b, we aligned 50-1000 sequences per genotype. For subtypes with fewer than 50 sequences in the GISAID database (40), we included all available sequences. For the H19 subtype, which is not yet added to the GISAID database, we aligned the three sequences published (41, 42). In the final step, we aligned the consensus cleavage loop sequences across all 19 subtypes. The resulting alignment (Fig. 6) shows that D346 is conserved in 13 of the 19 HA subtypes. It is substituted by alanine in H9, H12 and H19 HA, and by asparagine in H11, H13 and H16 HA. Quite intriguingly, H12, H13 and H16 HAs have previously been described as trypsin-resistant (8). X-ray crystallography revealed that H16 HA0 differs from its H1 and H3 counterparts by containing a short α-helix within the cleavage loop (21). To investigate whether N346 contributes to this resistance, we included H16 HA in our study and examined the effect of the inverse mutation, N346D, on HA0 cleavage efficiency.

**FIGURE 6.**
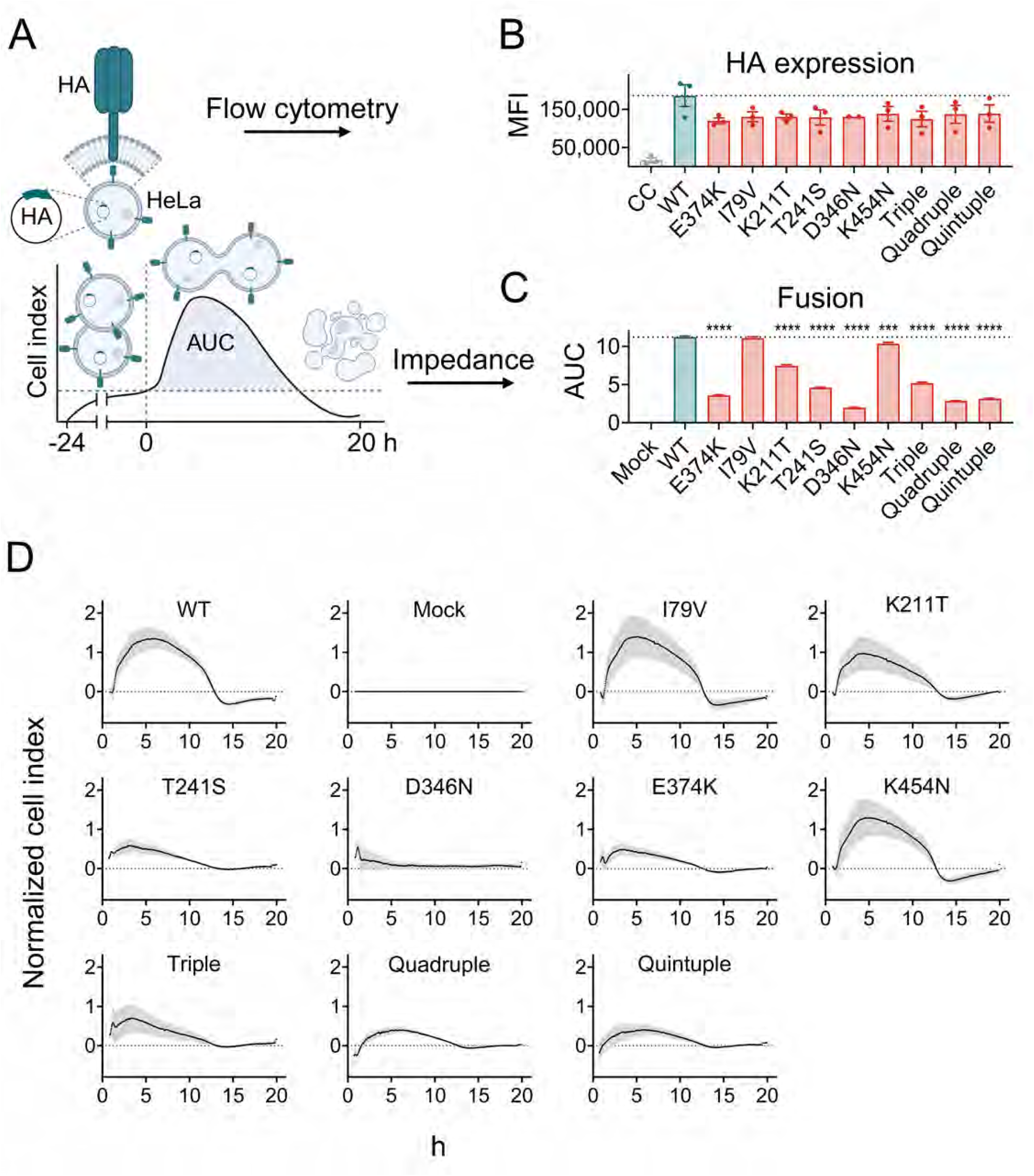
Cleavage loop alignment of H1-H19 HA0. Alignment of the cleavage loop sequences of each HA subtype. The scissile arginine is colored red and the residue at position 346 is boxed in teal. Dotted arrows indicated HA subtypes with A346 and full arrows indicated HA subtypes with N346.

### Mutation D346N impairs trypsin cleavage of H1 HA0, while N346D enhances cleavage of H16 HA0

To investigate whether the activation deficiency of the D346N-mutant H1 HA0 could be compensated by increasing the trypsin concentration, we generated H1N1 and H16N3 pseudoviruses bearing either WT or mutant HA (D346N for H1, N346D for H16). Following release from HEK293T producer cells, the pseudoviruses were incubated with a range of trypsin concentrations (i.e. 3.5-fold serial dilutions centered around 100 µg/mL), and subsequently analyzed for HA0 cleavage status and entry efficiency (experimental setup shown in Fig. 7A). Simple Western analysis (Fig. 7B) revealed that WT H1 HA0 was efficiently cleaved, with a detectable HA1 band already appearing at 2 µg/mL of trypsin, and approximately 60% of HA0 cleaved at 8 µg/mL. In contrast, the D346N mutant required ∼12-fold higher trypsin concentrations to reach comparable cleavage levels (note that the anti-H1 antibody detects only the HA1 cleavage product, while the anti-H16 antibody recognizes both HA1 and HA2). For H16 HA0, the N346D mutation had the opposite effect. The mutant proved more trypsin-sensitive than WT, with minimal cleavage seen at 8 µg/mL of trypsin compared to 28 µg/mL for WT. A direct comparison of the two WT proteins revealed that H16 HA0 is inherently less sensitive to trypsin than H1 HA0, requiring ∼100 µg/mL to achieve ∼50-60% cleavage, compared to ∼8 µg/mL for H1. These findings are consistent with previous reports (8, 21).

**FIGURE 7.**
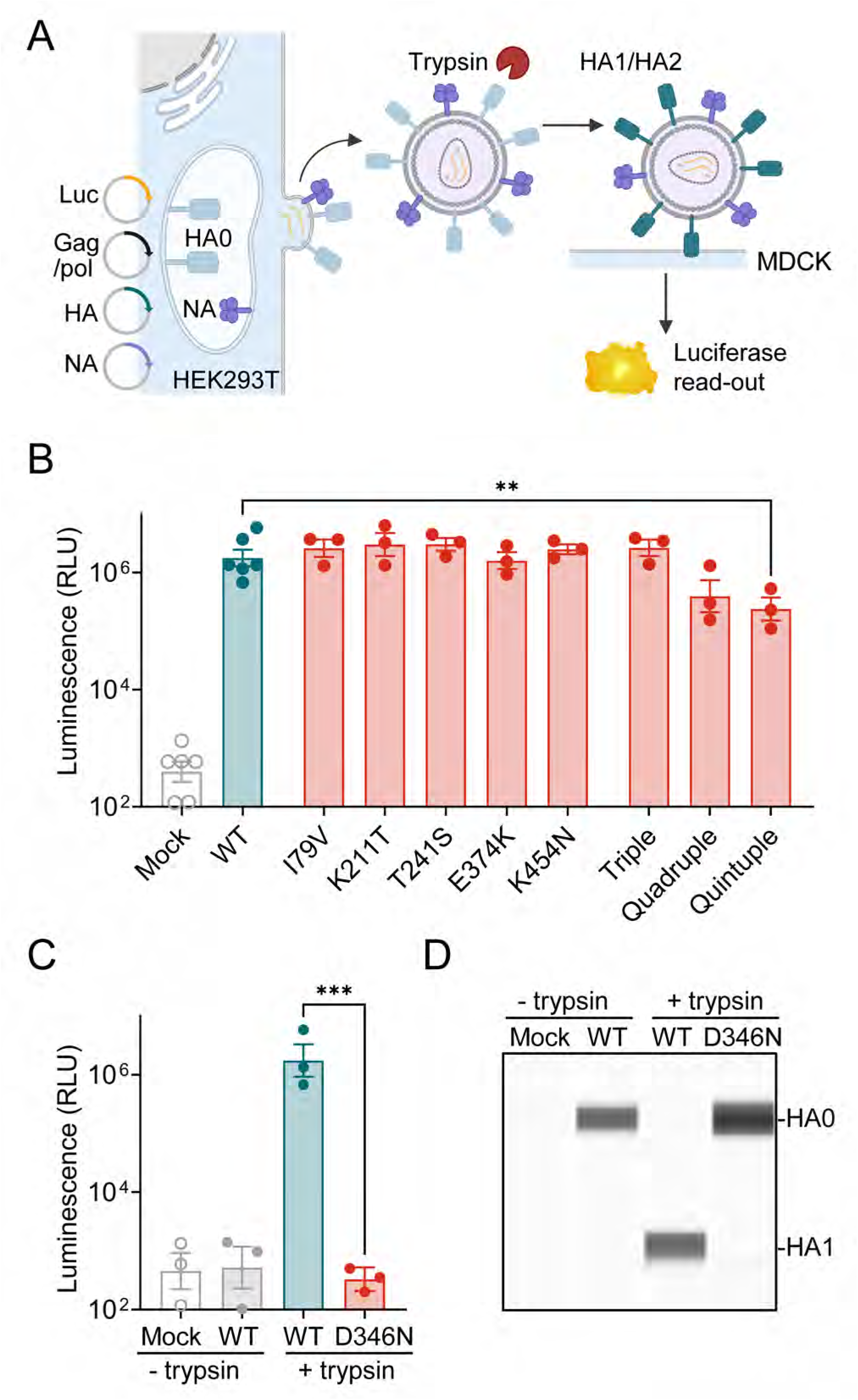
Trypsin-dependence of D346N-mutant H1 HA and the reverse mutant of H16 HA. (A) After production in HEK293T cells, the HA0-bearing pseudoviruses were exposed to different concentrations of trypsin. (B) Simple Western analysis of the HA0 cleavage status with antibodies recognizing the HA1 (for H1 HA) or HA1 and HA2 (for H16 HA) cleavage products. The images show a representative experiment, while the graphs show the mean values ± SEM (N=3) for the % of total HA that was present in cleaved form. (C) The pseudoviruses were incubated with MDCK cells for 24 h, followed by luciferase readout. Data points are the mean RLU values ± SEM (N=3). The dotted line represents the baseline value for pseudovirus that was not activated with trypsin. For each trypsin concentration, the difference between WT and mutant HA was analyzed by ordinary one-way ANOVA with Dunnett’s corrections for multiple comparisons.

In parallel, we conducted pseudovirus transduction assays in MDCK cells to assess viral entry efficiency, which closely mirrored the HA0 cleavage results (Fig. 7C). WT H1N1 pseudovirus exhibited maximum entry when activated by 8 μg/mL trypsin, while the D346N-mutant required 100 µg/mL. For H16N3, optimal entry was seen after incubation with 350 µg/mL trypsin for the WT and 100 µg/mL for the N346D mutant.

Taken together, mutation D346N impairs trypsin-mediated activation of H1 HA0, while the N346D mutation enhances cleavage of H16 HA0, partially compensating for the low activation efficiency of the WT. These reciprocal effects underscore the critical role of residue 346 in modulating HA0 activation by trypsin.

### Cleavage by TMPRSS2 is not significantly affected by mutating residue 346 and, in case of H16 HA, only occurs prior to virus release

While exogenous trypsin is commonly used to propagate IAV in cell lines lacking endogenous HA-activating proteases (e.g., MDCK cells) (43), viral activation in the human airways involves trypsin-like proteases such as transmembrane protease serine 2 (TMPRSS2), among others [reviewed in: (7, 44)]. TMPRSS2 is expressed both at the cell surface, with its protease domain facing the extracellular space, and within the trans-Golgi network (39), enabling it to cleave newly synthesized HA0 during intracellular trafficking (45). It remains unclear whether TMPRSS2 can also act extracellularly, after HA reaches the plasma membrane or the virus is released. To compare intracellular *versus* extracellular activation, we assessed pseudovirus activation by TMPRSS2 in two experimental setups (Fig. 8). In setup A, TMPRSS2 was co-expressed during pseudovirus production. Varying the amount of TMPRSS2 plasmid allowed us to modulate expression levels, as confirmed by flow cytometry (Fig. S2). In setup B, non-activated pseudoviruses were produced and subsequently used to transduce MDCK^TMPRSS2^ cells, which stably express abundant TMPRSS2 at the cell surface (Fig. S2).

**FIGURE 8.**
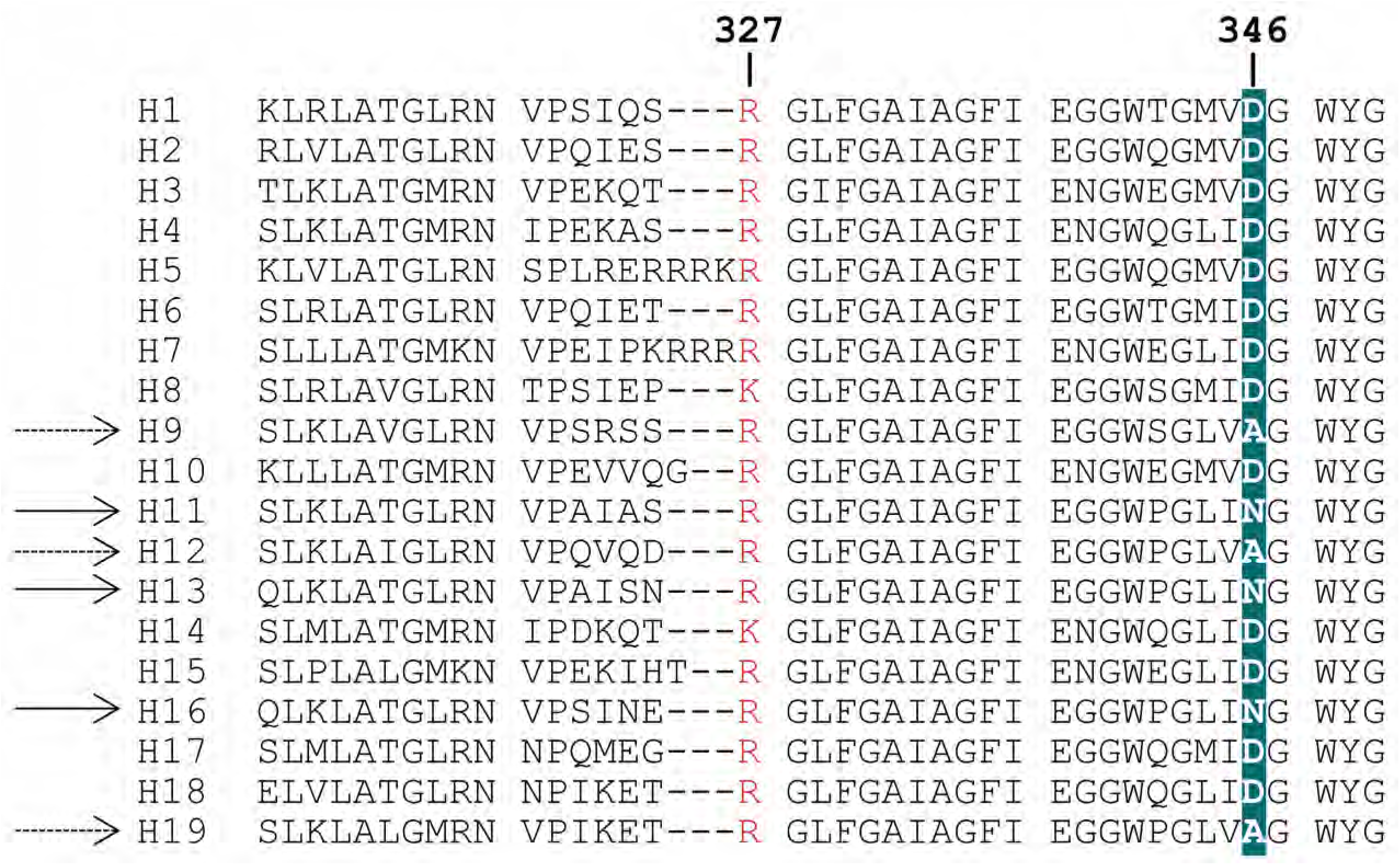
TMPRSS2-mediated activation of H16 HA0 depends on the subcellular location of this protease. (A) TMPRSS2-mediated HA0 cleavage was induced in the pseudovirus-producing cells, by co-transfecting a TMPRSS2-plasmid at different amounts. The activated particles were harvested and allowed to enter into MDCK cells. The graphs show the mean RLU values ± SEM (N=3) and the images show the HA0 cleavage status of pseudovirus generated using 35 ng of TMPRSS2 plasmid DNA per well. (B) Alternatively, the HA0-bearing pseudoviruses were harvested and exposed to TMPRSS2 during entry into MDCK^TMPRSS2^ cells. Individual and mean values ± SEM from 4 independent experiments. The difference between H1-WT vs. H16-WT; H1-WT vs. H1-mutant; and H16-WT vs. H16-mutant was analyzed by ordinary one-way ANOVA with Dunnett’s corrections for multiple comparisons.

Somewhat surprisingly, when TMPRSS2 was present in the producer cells (Fig. 8A), we saw no entry difference for the H1N1 pseudoviruses bearing WT or D346N-mutant HA. For H16N3, the N346D mutant showed slightly reduced activation compared to WT, but this difference was not statistically significant. When TMPRSS2 activation occurred at the point of viral entry (Fig. 8B), neither H1N1 nor H16N3 pseudoviruses showed differences between WT and mutant forms. However, in this setup, a striking difference in entry was observed between WT H16N3 and WT H1N1 pseudovirus, where the WT H16N3 resulted in ∼1,000-fold lower signal. This large difference was not observed in setup A, where WT H16N3 and WT H1N1 both induced luminescence signals of ∼10^5^-10^6^ RLU. This indicates that robust cleavage of H16 HA0 by TMPRSS2 [which was also described by others (12)] occurred only when HA was co-expressed with TMPRSS2. For H1N1, the difference between setup A and setup B was much smaller (i.e. ∼10-fold). This highlights a markedly reduced susceptibility of H16 HA0 to extracellular cleavage by TMPRSS2.

To further explore protease specificity, we co-transfected pseudovirus producer cells with plasmids encoding TMPRSS4 or TMPRSS13, two other members of the TMPRSS family known to activate HA0 under certain conditions (26, 46, 47). TMPRSS4 efficiently activated both H1 and H16 HAs (Fig. S3), regardless of whether the HA was WT or mutant. Although entry levels were significantly higher for H1N1 compared to H16N3 pseudovirus (*P* < 0.001), the latter generated robust signals (∼10^5^ RLU) when produced in the presence of TMPRSS4. In contrast, TMPRSS13 proved a very poor activator of H16 HA (both WT and mutant), while it activated H1 HA efficiently. Notably, activation of H1 HA by TMPRSS13 was significantly impaired by the D346N mutation (*P* < 0.001 versus WT). Given the limited research on TMPRSS13 in the context of IAV, this observation is difficult to interpret. One plausible explanation is that TMPRSS13 cleaves HA0 during virus budding, when HA0 is exposed at the plasma membrane, or extracellularly after viral release. This may explain the parallels observed with exogenous trypsin activation, which also acts extracellularly.

Taken together, the D346N-mutation in H1 HA0 caused a severe activation defect when exogenous trypsin was used, but had little to no effect when activation was mediated by cell-associated TMPRSS2 or TMPRSS4. We confirmed (8, 21) that H16 HA0 is relatively resistant to trypsin, and showed that this is partly alleviated by mutation N346D. On the other hand, TMPRSS2 smoothly activates H16 HA0, but only when present in the virus-producing cells. Once non-activated virus is released, the H16 HA0 protein faces poor cleavability by extracellular proteases.

### TMPRSS2 overcomes the trypsin resistance of H16 HA0 during virus rescue

The H16 HA sequence used in this study originates from an H16N3 virus isolated from gulls (48). Reverse engineering of this virus has proven challenging (49), likely due to the well-documented trypsin-resistance of H16 HA0 (8, 21). To investigate how residue 346 influences HA0 activation and virus rescue under different protease conditions, we reverse-engineered chimeric H1N1 (Virg09; WT and D346N) and H16N3 (Gull99; WT and N346D) viruses on a PR8 backbone. Two experimental setups were used to expose HA either to exogenous trypsin or to cell-associated TMPRSS2 (Table 1). In the first setup, the eight reverse genetics plasmids were transfected in a co-culture of HEK293T and MDCK cells, followed by virus expansion in MDCK cells, both in the presence of 5 µg/mL trypsin. Under these conditions, only the two Virg09_PR8_ viruses were successfully rescued, with the WT virus reaching titers approximately 100-fold higher than the D346N mutant (Table 1). In the second setup, a ninth plasmid encoding TMPRSS2 was included, and virus rescue was performed using a co-culture of HEK293T and MDCK^TMPRSS2^ cells for transfection, followed by virus expansion in MDCK^TMPRSS2^ cells. This approach enhanced virus yield for both Virg09_PR8_ viruses, particularly the D346N mutant, and was essential for rescuing the Gull99_PR8_ viruses, with the N346D mutant producing slightly higher titers than the WT virus. Sanger sequencing confirmed that neither of the four rescued viruses acquired additional mutations.

**TABLE 1.**
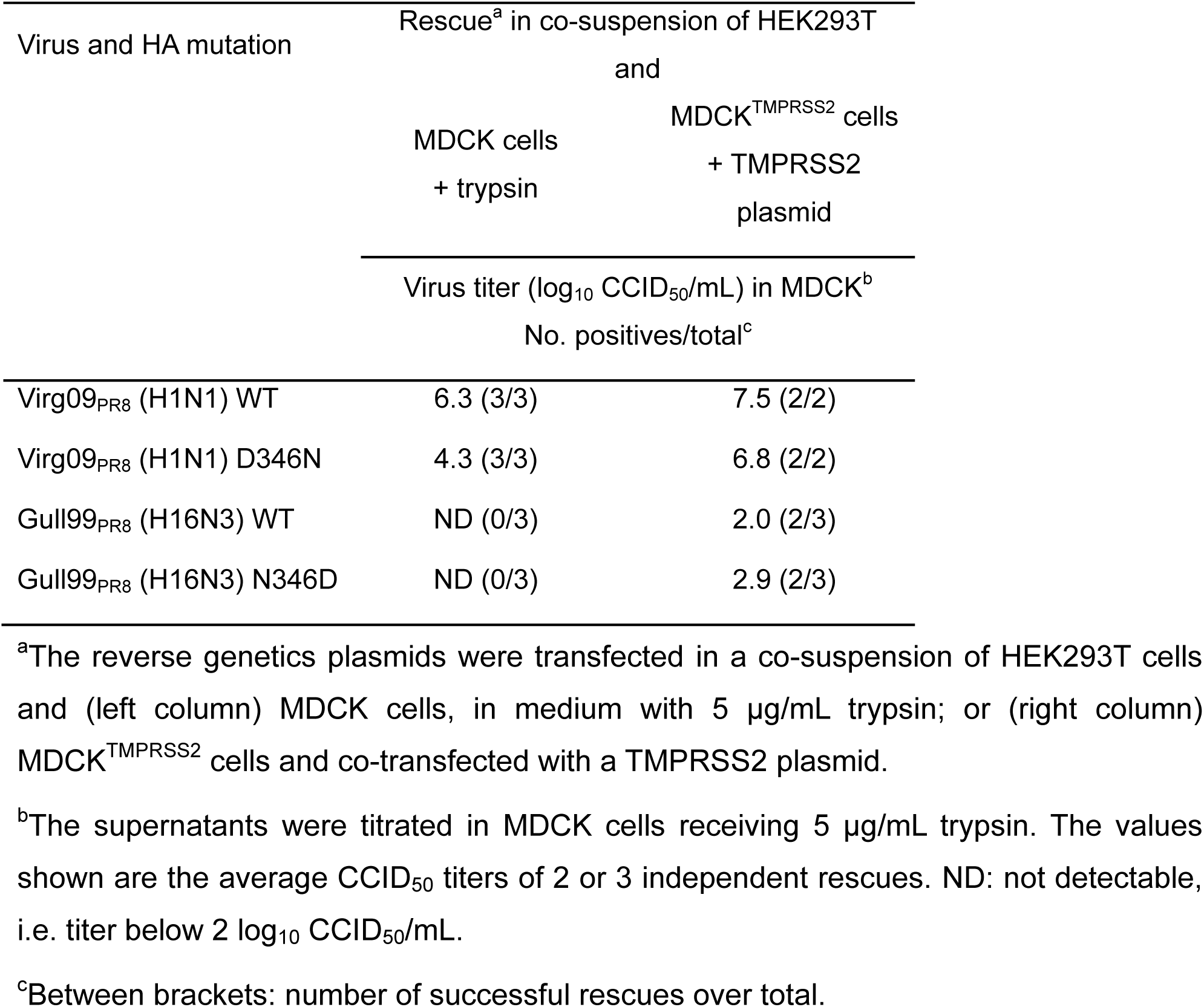
Virus rescue of Virg09_PR8_ (H1N1) and Gull99_PR8_ (H16N3) in MDCK and MDCK^TMPRSS2^ cells.

Growth kinetics further supported these findings. In MDCK cells treated with trypsin, the Virg09_PR8_-D346N virus replicated more slowly than the WT (*P* < 0.01 at 24 h p.i.; Fig. 9), whereas in MDCK^TMPRSS2^ cells, the difference was less pronounced (*P* < 0.05 at 24 h p.i.). For Gull99_PR8_, the N346D mutant replicated with similar kinetics as the WT in both cell types. However, sequencing of progeny viruses revealed that the Gull99_PR8_-WT virus acquired the N346D substitution by 72 h p.i. in both MDCK and MDCK^TMPRSS2^ cells (Fig. 9, insets). Analysis of the initial virus stock showed that a minor subpopulation had already acquired this mutation during virus expansion (Fig. 9, 0 h inset), further supporting the notion that D346 is strongly favored over N346 for efficient IAV replication in MDCK cells.

**FIGURE 9.**
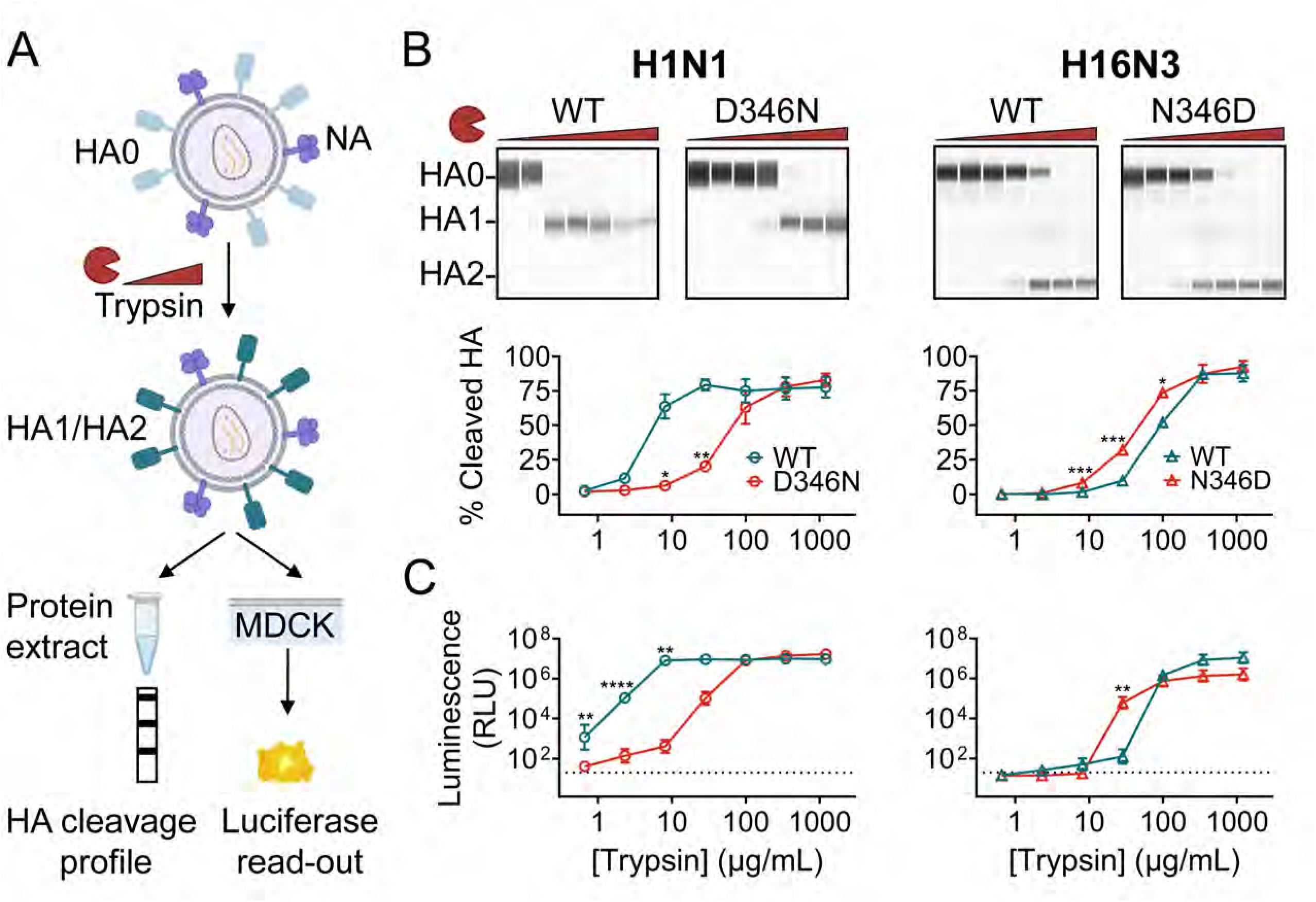
Growth kinetics of WT and mutant Virg09_PR8_ (H1N1) and Gull99_PR8_ (H16N3) viruses in MDCK and MDCK^TMPRSS2^ cells. MDCK and MDCK^TMPRSS2^ cells were infected with the indicated viruses at a multiplicity of infection (MOI) of 200 CCID_50_. Supernatants were collected at 24, 36, 48 and 72 h p.i., and the virus titers were determined in MDCK cells. Data points are the mean ± SEM (N=3). At each time point, statistical significance is shown for the difference between mutant and WT (multiple unpaired t-tests, with Holm-Šídák’s correction for multiple comparisons). Right to the H16N3 graphs, a zoom is shown of the HA sequence. Whereas the virus stock carried a mixed population of N346/D346, the D346 form dominated after 72 h replication in MDCK and MDCK^TMPRSS2^ cells.

### Besides its poor activation by exogenous trypsin, H16 HA is atypical in having an unusually low fusion pH

Given that a very high concentration of trypsin was required to activate the D346N-mutant form of H1 HA, we applied the same condition to determine the fusion pH in a cell-cell fusion assay. The luciferase-based method proved unsuitable, as the overlay cells did not tolerate high trypsin concentrations 24 hours after detachment. To overcome this, we employed a split-GFP assay (Fig. 10A) with two stable HeLa transfectants, expressing either the first ten beta sheets of GFP (GFP1-10), or the eleventh beta sheet (GFP11), each of which are non-fluorescent until they self-reassemble upon cell-cell fusion. A co-suspension of the two cell lines was transfected with HA plasmid, and two days later, the monolayers were treated with trypsin and exposed to a range of acidic buffers (pH 4.5-6.0). After 24 hours, green fluorescent syncytia were quantified by high-content imaging.

**FIGURE 10.**
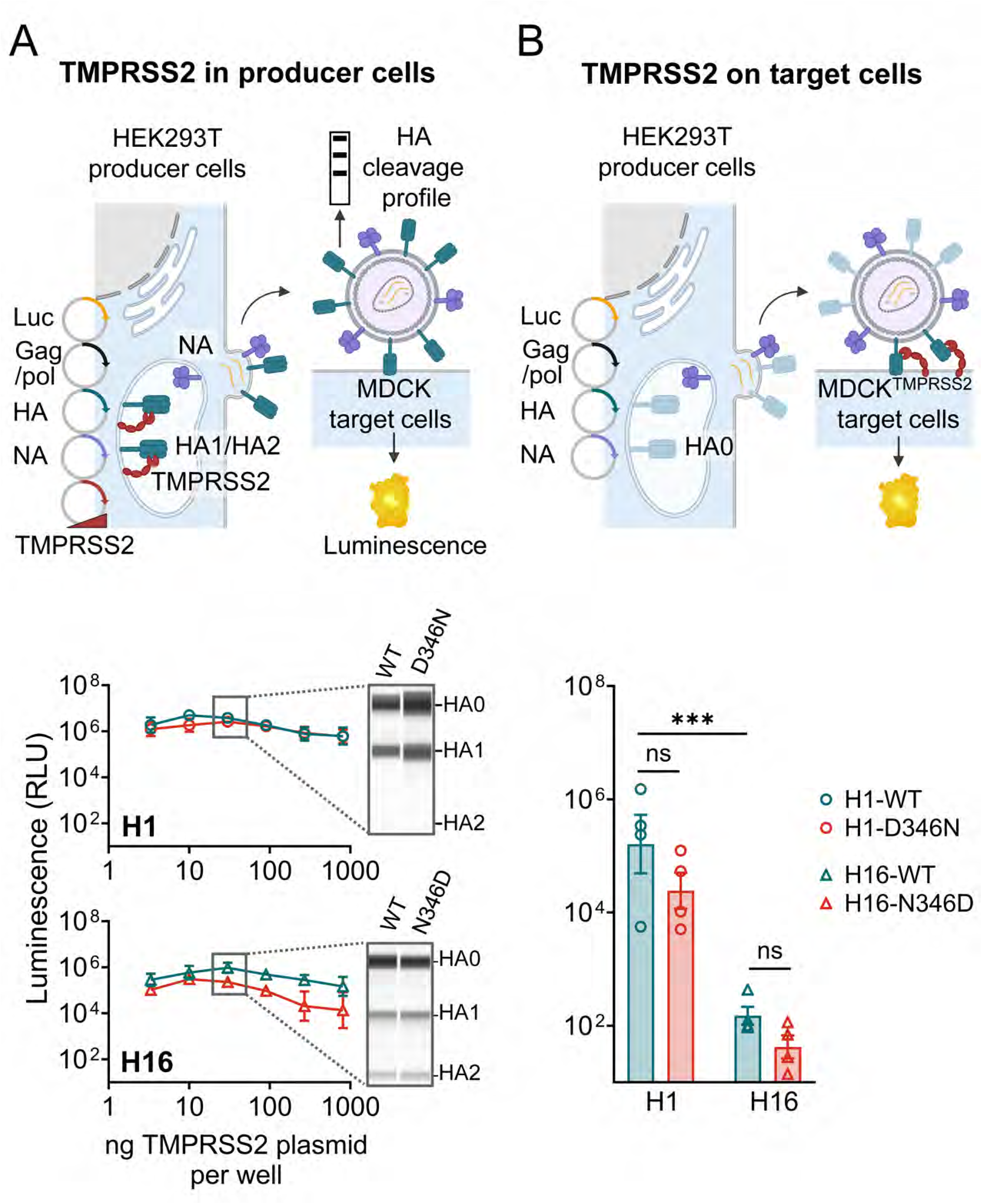
Fusion pH of D346N-mutant H1 HA and the reverse mutant of H16 HA. (A) Scheme of the split-GFP cell-cell fusion assay in HA-transfected cells exposed to low pH. (B) pH profiles for WT and D346N-mutant H1 HA (top), and WT and N346D-mutant H16 HA (bottom). The Y-axis shows the % cell-cell fusion, calculated by dividing the GFP area (after background subtraction) at each pH by the one at pH 5.0 (H1) or pH 4.5 (H16). The legend shows the fusion pH, defined as the pH where fusion was 50% relative to pH 5.0 (H1) or pH 4.5 (H16). Statistical significance is shown for the difference between mutant and WT (Extra sum of squares F test of best-fit value). Data points are the mean ± SEM (N=3). The insets show representative images of the green fluorescence in the condition exposed to pH 5.0 (H1) or pH 4.5 (H16) of each WT and mutant.

The results (Fig. 10B) show that the fusion pH was nearly identical (*P* > 0.05) between WT and D346N-mutant H1 HA, as well as between WT and N346D-mutant H16 HA (4.82 vs 4.89). The most striking observation was the unusually low fusion pH of 4.82 for H16 HA. Given that our H16 HA sequence is derived from a gull H16N3 isolate, we had expected a higher fusion pH in the range of 5.6-6.0, which is typical for avian HAs (50).

In short, though limited in scope, our experiments with replicating Gull99_PR8_ virus confirm the resistance of H16 HA0 to exogenous trypsin. A second peculiarity of H16 HA is its exceptionally low fusion pH of 4.8, far below the values typically observed for other avian HAs. This may reflect a host-specific adaptation, as H16N3 viruses have so far only been detected in gull species (51).

### Interpretation of the biological data in relation to the HA protein structure

With the exception of mutation D346N, the mutations identified in this study primarily have a stabilizing effect on H1 HA, since they reduce the fusion pH. To relate this biological effect to structural features of the protein, we mapped the mutated residues onto the crystal structure of prefusion A/Darwin/2001/2009 H1 HA [PDB ID 3m6s (52)], obtained at neutral pH, and onto cryo-EM models of the pre-fusion, post-fusion and intermediate states, published by Benton and colleagues (4). Our mutations are situated in the globular head as well as the stem domain (Fig. 11A), consistent with previous findings that stabilizing mutations can occur throughout the entire HA (53, 54).

**FIGURE 11.**
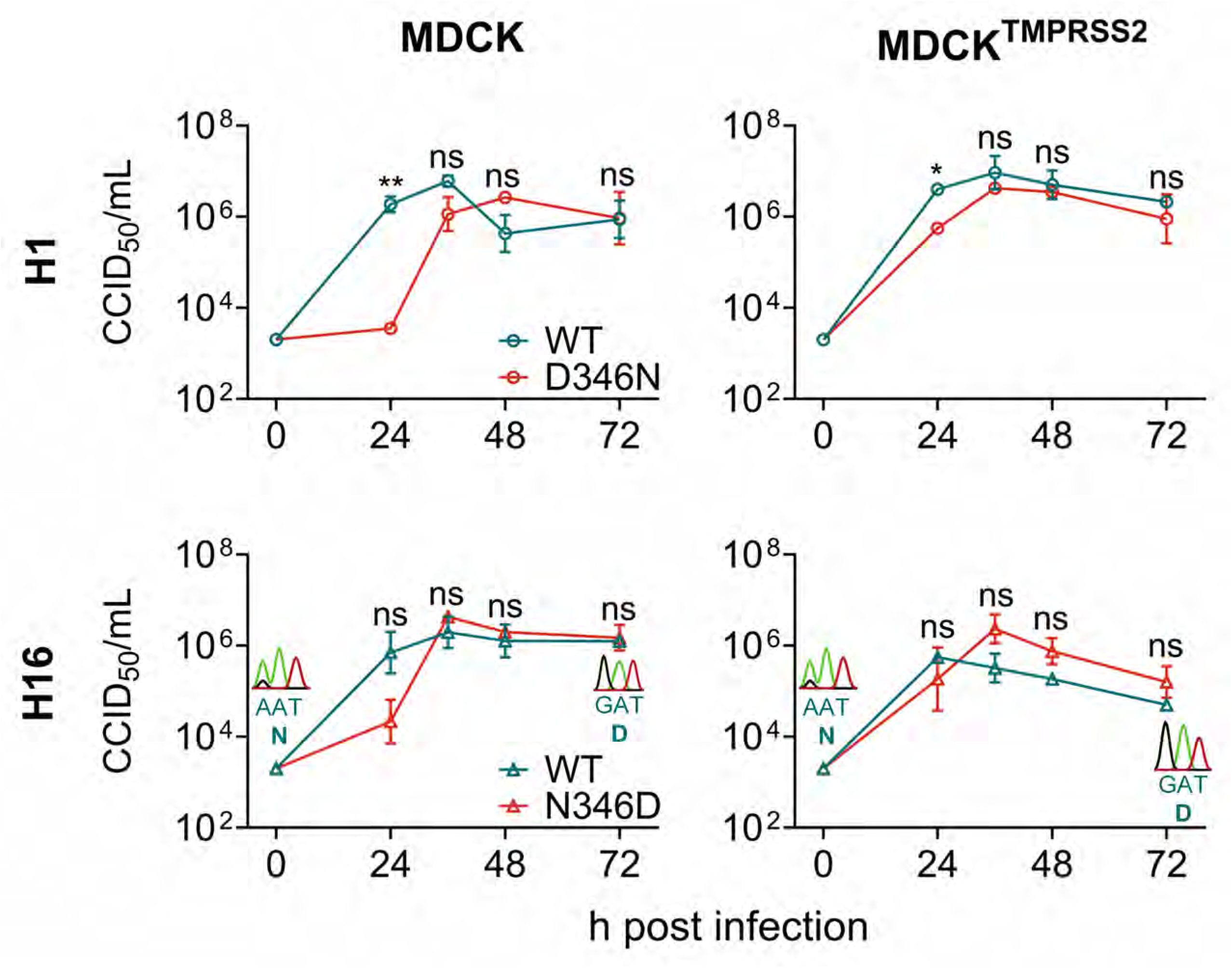
Location of the stability- and cleavability-determining residues in the HA protein structure. (A) Left panel: HA trimer in pre-fusion conformation, shown as white surface with one protomer colored according to receptor (R; in teal), vestigial esterase (E; in beige) and fusion (F; in purple) subdomain. The five sites at which we identified a stabilizing mutation are shown as red spheres. Right panel – upper figure: top view visualizing the location of K211 and T241 at an inter-protomer interface. Right panel – lower figure: I79 lies in a hydrophobic patch adjacent to the 110-helix of the E subdomain, which interacts with the interhelical loop of the F subdomain. (B) The low pH-induced transition from the pre-to post-fusion structure starts with dilation of the three head domains. After formation of the extended coiled-coil (shown in pink, green, orange and blue in the extended intermediate), the C-terminal part (colored in blue with K454 shown as red sphere) folds back by 180°, resulting in the final post-fusion structure. Models generated from PDB 3M6S (with few residues altered to fit Virg09) and PDB 6Y5K and PDB 1HTM (using SWISSMODEL to model Virg09). (C) Model of the HA0 precursor protein with the location of the cleavage loop marked by a black box. (D) In the X-ray structure of the H16 HA0 cleavage loop (PDB 4F23) (21), the scissile R327 residue (in red) is hidden behind a short α-helix (in purple). Residue N346 is shown in teal. (E) Based on an Alphafold model for H1 HA0, mutation D346N (in teal) is predicted to induce an extra turn (in purple) in the cleavage loop, which may reduce the access of R327 (in red) to proteases.

Although the most drastic refolding occurs in the stem, the HA head is the first region to respond to acidification. Dilation of the three head domains is required to expose the interhelical loop in the F subdomain. This offers a possible explanation for the stabilizing effect of mutations K211M and T241S, which are located at an inter-protomer interface in the R subdomain (Fig. 11A; top view on the right). K211M causes a loss of net charge, while maintaining sidechain flexibility. Since K211T was found to reduce the fusion pH, we presume that the positive charge at this position may play a role in the initial stages of trimer opening at low pH. T241S reduces sidechain size while preserving polarity. Although structural analysis at neutral pH did not reveal direct interactions between K211 or T241 and residues of neighboring head domains, our biological data suggest that these residues may influence head domain rearrangement at acidic pH.

Residue I79 is located in the E subdomain, which connects the R and F subdomains. Intriguingly, it is located in a hydrophobic patch adjacent to the 110-helix of the R subdomain (Fig. 11A; bottom right). This helix interacts with the interhelical loop of the F subdomain, which undergoes a loop-to-helix transition at low pH. Given that mutations in the 110-helix have been shown to affect acid-stability of H5 HA (55), the same seems valid for mutations in the nearby hydrophobic patch. The I79V mutation retains sidechain hydrophobicity but slightly reduces its size.

Once the interhelical loop of the F subdomain transitions into a helix, a long coiled-coil structure is formed that extends towards the head domain (Fig. 11B). Then, the C-terminal part of the long helix folds back by 180°, causing its relocation upwards and alongside the coiled-coil core of the final post-fusion structure. Residue K454 is situated in this fold-back region (Fig. 11B: structure on the right). Mutation K454N causes a loss of net charge, besides reducing the size and flexibility of the sidechain. Each of these factors may have an effect on the local rearrangements in this part of the HA stem. As a reference, we included the known stabilizing mutation E374K, which is located in the short helix of the F subdomain and introduces a salt bridge between K374 and E21 of the adjacent monomer, thereby enhancing H1 HA trimer stability (17, 19).

To understand why mutation D346N impairs trypsin-mediated cleavage of H1 HA0, we used an entirely different approach. D346 lies within the cleavage loop, part of which becomes the fusion peptide upon cleavage. The few HA0 subtype structures that are so far available show extensive differences in the folding of the cleavage loop. This part of the protein is very flexible, which may influence the accessibility of the cleavage site and consequently its susceptibility to proteases. In the crystal structure of Gull99-H16 HA0, a short helix positions residue R327 deep in the stem domain (Fig. 11D), likely contributing to its resistance to trypsin cleavage (21).

We used AlphaFold to generate models of Virg09 HA0 and study its cleavage loop in both WT and D346N mutant forms. Interestingly, the two models predicted distinct conformations near the mutation site (Fig. 11E). In the WT model, the loop is more solvent-exposed, whereas in the D346N model, an additional turn buries the loop deeper into the HA stem. Consequently, the scissile R327 residue is accessible in the WT but less so in the D346N mutant, offering a structural explanation for the observed cleavage defect.

## DISCUSSION

The influenza virus HA is a highly dynamic and adaptable protein that is subject to structural instability, antigenic drift and host adaptation. Understanding how these processes are modulated at the level of specific HA regions or amino acid residues remains a key area of interest. In this study, we employed random mutagenesis, a broad and unbiased approach, to identify mutations that increase the acid-stability of HA. While previous HA random mutagenesis studies have focused on receptor binding affinity (56) or overall viral replication fitness (22, 57, 58), our work specifically targeted mutations that enhance HA acid-stability, given its relevance to influenza virus transmission, host adaptation (16, 18, 50, 59) and vaccine stability (17, 19, 20, 60–62). We considered the HA of an early A(H1N1)pdm09 virus to be a relevant case, given its rather weak stability, which increased *via* the acquisition of stabilizing mutations upon sustained circulation in the human population (16–18).

By combining random mutagenesis with selection of mutant viruses that survive incubation at pH 5.0, we identified four novel sites (i.e. I79, K211, T241 and K454) across the H1 HA protein that are amenable to stabilizing mutations. While there is already extensive literature on HA mutations that alter the acid-stability of H3, H5 and H7 subtypes, data for H1 HA remain more limited (17, 54). Moreover, most previously reported mutations tend to increase, rather than decrease, the fusion pH. Although we initially expected a bias towards mutations in the HA stem, where several pH-sensitive residues can trigger refolding of HA via electrostatic effects (62), three out of four stabilizing mutations that we selected were located outside the stem domain. Our expectation was supported by the well-characterized stem mutation E374K, which we included as a reference. The strong stabilizing effect of K374 has been attributed to the formation of an inter-monomeric salt bridge that reinforces trimer integrity (17, 19, 63). Nonetheless, our findings align with previous studies (17, 18) that mutations in the globular head, vestigial esterase and membrane-proximal part can have a profound impact on HA acid-stability, particularly when located at inter-monomer interfaces or regions critical for low-pH induced structural transitions. We also observed that combining multiple stabilizing mutations had no stronger effect on the fusion pH than the individual mutations, since the combinations that we tested were, at best, modestly additive. This conclusion was also made in a previous study (37). The authors proposed a hierarchical influence of mutation sites, where substitutions in the short α-helix of the stem domain may override those in the membrane-distal region, and substitutions near the fusion peptide may dominate over those in the other two regions. Translated to our study, this would mean that the E374K mutation (in the short α-helix) overrides the effects of mutations in the head (K211T and T241S) and vestigial esterase (I79V) domains. Anyway, the possibility that stabilizing HA mutations can occur in diverse regions of the protein complicates the interpretation of HA sequence changes during zoonotic IAV surveillance.

In retrospect, using more stringent selection conditions (e.g. lower pH or longer incubation) might have yielded mutants with even greater acid-stability. However, an inactivation pH of 4.9-5.0 seems to be the lower threshold tolerated by replication-competent viruses, with only a few strains (such as the lab-adapted PR8 strain) known to exhibit such prominent stability (59, 64). While engineered mutations at key residues in the stem domain can yield HA proteins stable at pH 4.7 (62) or even 4.3 (65), such extreme stabilization likely impairs viral replication. After endocytosis, influenza virus particles encounter progressively acidic environments: early endosomes (pH 6.5-6.0), late endosomes (pH 5.5-5.0) and lysosomes ( pH 4.5-5.0) (66–68). Hence, a virus with a very low fusion pH may fail to fuse before reaching lysosomes, where it faces interferon-inducible factors (69, 70). In this regard, our observation that the Gull99-H16 HA has a fusion pH of ∼4.8 in HA-transfected HeLa cells is particularly intriguing. This value is unusually low for an avian HA and reminiscent of the remarkably low HA activation pH (i.e. 5.0) reported for an avian H1N1 virus from shorebirds (71). To our knowledge, this is the first report highlighting the distinctively low fusion pH of H16 HA. Previous studies either failed to determine its fusion pH, likely due to insufficient trypsin activation (8), or reported a much higher value of 5.5 using a syncytium assay with live Gull99 H16N3 virus and Vero cells (64). Follow-up studies may help to resolve this discrepancy.

Our study also addressed whether altered HA fusion properties have an impact on viral replication fitness. Besides rendering H1 HA more acid-stable, all five HA mutations that we studied impaired cell-cell fusion efficiency, as evidenced by smaller syncytia in pH 5-treated HA-expressing cells. Our impedance-based fusion assay proved highly suitable for real-time monitoring of cell-cell fusion, yielding additional information on the biological effect of HA mutations that complements assessment of the fusion pH. Despite reducing the fusion-promoting activity of HA, none of the mutations altered pseudovirus entry efficiency into MDCK cells or replication of reverse-engineered viruses. The mutant viruses reached titers comparable to WT and the introduced mutations remained stable over four serial passages. This holds even true for mutation E374K, which combines a fusion pH as low as 5.17 with a pronounced reduction in syncytium size. This may be due to the low endosomal pH of MDCK cells, which can support the replication of acid-stable viruses (19). In line with our findings, another recent mutagenesis study also found no strong correlation between HA acid-stability and viral entry efficiency (72). The smooth replication of HA-stabilized viruses is important for vaccine production, where mutations at key sites in the HA trimer may help to produce a live attenuated or inactivated vaccine with superior shelf-life, without compromising viral yield.

In addition to the stabilizing mutations, we identified another HA mutation, D346N, which severely impaired trypsin-mediated HA activation. The survival of D346N-mutant H1N1 virus during the pH 5.0 selection likely reflects a much higher proportion of uncleaved, acid-stable HA0 on the viral surface. D346N-mutant H1N1 pseudovirus required 12-fold more trypsin for activation than WT, while the reciprocal N346D-mutation in H16N3 pseudovirus reduced the trypsin requirement by 4-fold. These findings suggest that the presence of N346 alters the conformation of the cleavage loop, reducing its accessibility to trypsin. Validation of this hypothesis will require structural investigation.

A somewhat puzzling observation was the location-dependent activity of TMPRSS2. We confirmed two other reports that H16 HA0 is readily activated by TMPRSS2, which is in sharp contrast to its trypsin-resistance (8, 12). However, this activation occurred only when TMPRSS2 was expressed in the same cells responsible for HA synthesis, processing and membrane trafficking. This aligns with earlier findings that TMPRSS2 primarily cleaves H1 HA0 intracellularly rather than at the cell surface (45). In our experiments, this preference for intracellular processing was evident for both H1 and H16 HAs, though it was more pronounced for the H16 subtype. Together, our trypsin and TMPRSS2 data establish that, unlike most HA subtypes, H16N3 virus is barely activated by extracellular proteases. In contrast, when TMPRSS2 is present in the virus-producing cells, H16 HA0 is cleaved nearly as efficiently as H1 HA0, with minimal differences observed between WT and D346N/N346D mutants. This suggests that, especially for H16 HA0, the shape and accessibility of the cleavage loop profoundly change during HA biosynthesis, glycosylation, trimerization, trafficking to the plasma membrane or viral budding and release. It is plausible that the short α-helix shielding the scissile arginine in H16 HA0 only forms at a later stage (21, 73–75). This requirement for cleavage within virus-producing cells was evident not only in the pseudovirus experiments but also during reverse genetics, where co-expression of TMPRSS2 was essential to rescue the Gull99_PR8_ virus. This aligns with a previous report where H16N3 virus could not be rescued in trypsin-supplemented HEK293T/MDCK cells, but was successfully recovered by inoculating transfected HEK293T cells into embryonated hen eggs (49). Once rescued, the virus showed normal growth kinetics in trypsin-supplemented medium, likely due to prolonged exposure (three days), compared to the brief trypsin treatment (15 min) during pseudovirus production. Similarly, the D346N-mutant Virg09_PR8_ virus showed only a transient growth delay relative to WT.

We would like to note that, in our experiments with H16N3 (pseudo)virus, we primarily employed trypsin and TMPRSS2 as model proteases to facilitate comparison with H1N1 and to compare extra-*versus* intracellular cleavage. The identity of the proteases responsible for activating H16N3 virus in its natural host, gulls, remains unknown (49). Still, our findings could be relevant for the potential role of this gull virus in influenza virus ecology and animal-to-human spillover (76, 77). First, the low fusion pH (∼4.8) of H16N3 resembles that of human IAVs which typically have lower fusion pH values than avian strains (16). Combined with its prominent ability to bind α2,6-linked sialylated glycans (78) and undergo intracellular activation by human TMPRSS2 [this study and (8, 12)], the gull H16N3 virus appears well-adapted for attachment to the human respiratory tract [as observed in (79)] and for replication within this tissue. Second, its high acid-stability and resistance to cleavage by extracellular proteases may enhance environmental persistence. Gulls inhabit coastal areas where the high salinity of sea water can reduce viral persistence (80) and promote inactivation in evaporating droplets (81, 82). The H16N3 virus may be well-adapted to survive under these harsh circumstances.

To conclude, our random mutagenesis and pH-based selection approach revealed four novel stabilizing mutations in H1 HA, located across multiple domains. This highlights the complex interplay between all parts of HA during low pH-induced refolding. Although these mutations reduced the fusion-promoting activity of HA, they did not impair viral entry or replication. A fifth mutation, D346N, drastically reduced trypsin-mediated cleavage but not intracellular activation by TMPRSS2. The presence of N346 in gull-derived H16 HA, which we found to exhibit an unusually low fusion pH, contributes to its trypsin-resistance. Together, our results yielded new insights into the determinants of HA stability and cleavability, with implications for IAV surveillance and vaccine production.

## SUPPLEMENTAL FIGURES

**FIGURE S1. pH profiles of other mutations identified by random mutagenesis of H1 HA.** pH profiles obtained from cell-cell fusion assays with WT HA (Virg09_ecto_; black curve that is shown in each panel); and five mutants (curves shown in red). The Y-axis shows the % cell-cell fusion relative to the pH 5.0 condition, calculated by dividing the luminescence signal at each pH by the one at pH 5.0 after background subtraction. The legend shows the fusion pH, defined as the pH where fusion was 50% relative to pH 5.0. Statistical significance is shown for the difference in fusion pH between mutant and WT (Extra sum of squares F test of best-fit value). The data points are the mean ± SEM (N=3-4).

**FIGURE S2. Flow cytometry to verify robust and plasmid concentration-dependent TMPRSS2 expression.** The data show the TMPRSS2-positivity of the cell population, assessed in either HEK293T producer cells transfected with different amounts of TMPRSS2 plasmid (xy graph and scatterplot on top) or the stably transduced MDCK^TMPRSS2^ target cells (bar graph and scatterplots on the bottom). The xy graphs show the mean ± SEM (N=3). In the bar graphs, each data point is shown and the bar represents the mean. The scatterplots are taken from one representative experiment.

**FIGURE S3. Entry of pseudovirus activated by TMPRSS4 or TMPRSS13 during pseudovirus production.** (A) Setup of the luciferase based pseudovirus entry assay. TMPRSS4 or TMPRSS13 plasmid was added to the transfection mix to activate the pseudovirus during production. The virus was then allowed to enter in MDCK cells, followed by luciferase readout after 72 h. (B) RLU values for the different pseudoviruses. Data points are the mean ± SEM (N=3). Statistical significance was analyzed by ordinary two-way Anova with Tukey correction for multiple comparisons.

## ACKNOWLEDGEMENTS

We gratefully acknowledge all data contributors, i.e., the Authors and their Originating laboratories responsible for obtaining the specimens, and their Submitting laboratories for generating the genetic sequence and metadata and sharing via the GISAID Initiative, on which this research is based.

Molecular graphics and analyses were performed with UCSF ChimeraX, developed by the Resource for Biocomputing, Visualization, and Informatics at the University of California, San Francisco, with support from National Institutes of Health R01-GM129325 and the Office of Cyber Infrastructure and Computational Biology, National Institute of Allergy and Infectious Diseases.

S.R. is holder of an SB-PhD fellowship from the FWO Research Foundation Flanders (No. 1S92321N). The authors wish to thank Marie Horemans for helping with the Nanopore sequencing; Joost Schepers for his help with quantification of the high-content imaging data; S. Pöhlmann for the generous donation of plasmid materials; and M. Matrosovich and J. Bloom for providing cell lines.

